# Single phage proteins sequester TIR- and cGAS-generated signaling molecules

**DOI:** 10.1101/2023.11.15.567273

**Authors:** Dong Li, Yu Xiao, Weijia Xiong, Iana Fedorova, Yu Wang, Xi Liu, Erin Huiting, Jie Ren, Zirui Gao, Xingyu Zhao, Xueli Cao, Yi Zhang, Joseph Bondy-Denomy, Yue Feng

**Affiliations:** Beijing Advanced Innovation Center for Soft Matter Science and Engineering, State Key Laboratory of Chemical Resource Engineering, College of Life Science and Technology, Beijing University of Chemical Technology, Beijing 100029, China; Ministry of Education Key Laboratory of Protein Science, Beijing Advanced Innovation Center for Structural Biology, Beijing Frontier Research Center for Biological Structure, School of Life Sciences, Tsinghua University, Tsinghua-Peking Center for Life Sciences, Beijing 100084, China; Department of Microbiology and Immunology, University of California, San Francisco, San Francisco, CA, 94158, USA; State Key Laboratory for Biology of Plant Diseases and Insect Pests, Ministry of Agriculture, Institute of Plant Protection, Chinese Academy of Agricultural Sciences, Beijing 100081, China; Quantitative Biosciences Institute, University of California, San Francisco, San Francisco, CA 94158, USA; Innovative Genomics Institute, Berkeley, CA 94720, USA

## Abstract

Prokaryotic anti-phage immune systems use TIR (toll/interleukin-1 receptor) and cGAS (cyclic GMP-AMP synthase) enzymes to produce 1’’-3’/1’’-2’ glycocyclic ADPR (gcADPR) and cyclid di-/tri-nucleotides (CDNs and CTNs) signaling molecules that limit phage replication, respectively ^1–3^. However, how phages neutralize these common systems is largely unknown. Here, we show that <u>T</u>hoeris <u>a</u>nti-<u>d</u>efense proteins Tad1 ^4^ and Tad2 ^5^ both have anti-CBASS activity by simultaneously sequestering CBASS cyclic oligonucleotides. Strikingly, apart from binding Thoeris signals 1’’-3’ and 1’’-2’ gcADPR, Tad1 also binds numerous CBASS CDNs/CTNs with high affinity, inhibiting CBASS systems using these molecules *in vivo* and *in vitro.* The hexameric Tad1 has six binding sites for CDNs or gcADPR, which are independent from two high affinity binding sites for CTNs. Tad2 also sequesters various CDNs in addition to gcADPR molecules, inhibiting CBASS systems using these CDNs. However, the binding pockets for CDNs and gcADPR are different in Tad2, whereby a tetramer can bind two CDNs and two gcADPR molecules simultaneously. Taken together, Tad1 and Tad2 are both two-pronged inhibitors that, alongside anti-CBASS protein 2, establish a paradigm of phage proteins that flexibly sequester a remarkable breadth of cyclic nucleotides involved in TIR- and cGAS-based anti-phage immunity.

## Introduction

Bacteria encode numerous immune systems that protect them from phage infection ^6–11^. In turn, phages also developed mechanisms to antagonize these immune systems and effectively replicate, such as expressing proteins with anti-immune activities, out of which anti-CRISPR (Acr) proteins have been studied extensively ^12–15^. Up until now, phage anti-immune proteins have been discovered for many different systems, including CRISPR-Cas, restriction-modification, and BREX, which largely rely on protein-protein interactions to block immune function ^16^. However, recently discovered inhibitors of cyclic nucleotide-based anti-phage systems, like CBASS, Thoeris, Pycsar, and Type III CRISPR-Cas, have revealed the ability of phage proteins to sequester or degrade cyclic nucleotides ^4,5,17–20^.

The Thoeris anti-phage system encodes ThsB, a protein with a Toll/interleukin-1 receptor (TIR) domain, which senses phage infection and produces the 1”–3’ gcADPR signaling molecule that subsequently activates the NADase effector ThsA ^1,2,4^. CBASS (cyclic-oligonucleotide-based anti-phage signaling system) encodes a cGAS/DncV-like nucleotidyltransferase (CD-NTase) that produces cyclic dinucleotides (CDNs) or cyclic trinucleotides (CTNs) upon phage infection ^3^. A broad diversity of CD-NTases has been identified in bacteria ^21^, which are able to produce at least 12 different cyclic oligonucleotide species ^21–26^. These cyclic oligonucleotides also bind to a cognate effector, which is proposed to kill the cell and stop successful phage replication.

<u>T</u>hoeris <u>a</u>nti-<u>d</u>efense proteins Tad1 and Tad2 antagonize immunity by sequestering the signaling molecule 1”-3’ gcADPR as a sponge protein ^4,5^. For CBASS, two phage proteins have been discovered that antagonize its immunity. Anti-<u>cb</u>ass protein 1 (Acb1) degrades the cyclic oligonucleotide signals ^19^, and Acb2 is a sponge for CDNs ^17,18^ and CTNs at a distinct binding site ^27^. Here, we report the surprising observation that Tad1 and Tad2 both also possess anti-CBASS activity by sequestering a breadth of CBASS signals. Strikingly, apart from 1’’-3’ and 1’’-2’ gcADPR, Tad1 also binds to CBASS CDNs 2’,3’-/3’,2’-/3’,3’-cGAMP/cUA/cAA/cGG and CTNs cA_3_/3’3’3’-cAAG (cAAG) with high affinity. Tad2 sequesters CBASS CDNs 3’,3’/3’,2’/2’,3’-cGAMP/cGG/cUG in addition to gcADPR molecules. CBASS is generally more common than Thoeris ^28^, thus these findings greatly broaden our appreciation of the utility of these proteins to the many phages that encode them. Some bacterial species also encode both Thoeris and CBASS immune systems ^11,29^, which Tad1 and Tad2 could inhibit simultaneously. Tad1 and Tad2 are therefore two-pronged inhibitors that block Thoeris and CBASS activity due to the similar nature of their immune signaling molecules despite the independent evolutionary origins of the enzymes that create them.

## Results

### Tad1 sequesters CBASS cyclic trinucleotides and dinucleotides

Due to the overall similarities between signaling molecules used by multiple defense systems, we asked whether anti-Thoeris Tad1 and Tad2 sponges also sequester cyclic nucleotides used in Pycsar, CBASS, or Type III CRISPR-Cas signaling systems ^30–32^. Four cyclic mononucleotides (cCMP, cUMP, cGMP and cAMP), eight CDNs (3’,3’-cGAMP, cGG, cUG, cUA, cUU, cAA, 2’,3’- and 3’,2’-cGAMP), two CTNs (cA_3_ and cAAG), as well as cA_4_ and cA_6_ were tested. Surprisingly, native gel assays showed a shift of both CbTad1 (from *Clostridium botulinum* prophage) and CmTad1 (from *Clostridioides mangenotii* prophage) upon adding cA_3_ or cAAG, and a shift of CbTad1 upon adding 2’,3’-cGAMP or 3’,2’-cGAMP (Extended Data Figure 1). These binding events were further verified by isothermal calorimetry (ITC) experiments (Figure 1a, Extended Data Figure 2), which showed that CbTad1 and CmTad1 bind cA_3_ with a *K_D_* of ∼14.0 and 9.8 nM, respectively, and they bind cAAG with a *K_D_* of ∼12.5 and 20.1 nM, respectively (Figure 1a). CbTad1 bound 2’,3’-cGAMP and 3’,2’-cGAMP with a *K_D_* of ∼31.1 and 24.5 nM, respectively (Figure 1a, Extended Data Figure 2), however, only bound weakly (*K*_D_ >0.4 µM) to 3’,3’-cGAMP/cGG/cUA/cAA (Figure 1a, Extended Data Figure 2). High-performance liquid chromatography (HPLC) revealed that incubating CbTad1 with cA_3_ or 2’,3’-cGAMP depleted detectable molecules, but they were detected again in unmodified form after proteolysis of CbTad1 (Figure 1b). The strong binding values reported here are >1 order of magnitude stronger than the reported CmTad1 binding affinity for 1”-2’ gcADPR of 241 nM ^4^, which was the first identified ligand for this protein. Bioinformatic analysis of the *Clostridium* genus revealed CBASS CD-NTases that produce cA_3_/cAAG (CdnD) and 3’,2’-cGAMP (CdnG) (Extended Data Figure 3a) ^21,22^, highlighting the likely biological driver for the observed binding spectra of CbTad1 and CmTad1. Taken together, these results demonstrate that Tad1 also binds to and sequesters both CTNs and CDNs used in CBASS immunity in addition to gcADPR molecules.

**Figure 1.**
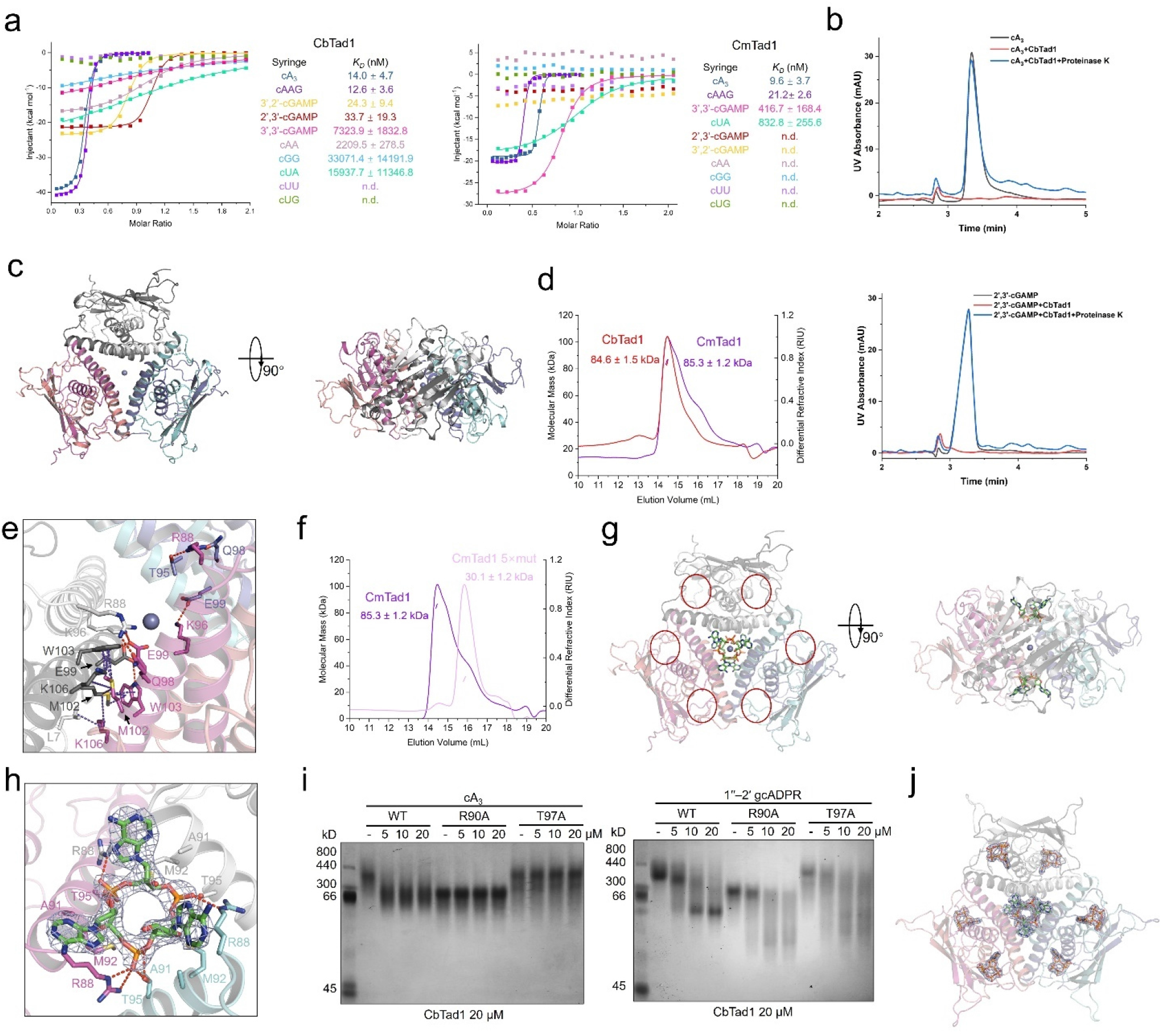
Tad1 is a hexamer to bind to two molecules of cyclic trinucleotides. **a**, ITC assays to test binding of cyclic oligonucleotides to CbTad1 and CmTad1. Representative binding curves and binding affinities are shown. The *K_D_* values are mean ± s.d. (n=3). Raw data for these curves are shown in Extended Data Figure 2. **b,** The ability of CbTad1 to bind and release cA_3_ and 2’,3’-cGAMP when treated with proteinase K was analyzed by HPLC. cA_3_ and 2’,3’-cGAMP standard was used as controls. The remaining nucleotides after incubation with CbTad1 were tested. **c,** Overall structure of CmTad1 hexamer. The Zn ion is shown as a sphere. Three views are shown. **d,** Static light scattering (SLS) studies of purified CbTad1 and CmTad1. Calculated molecular weight is shown above the peaks. **e,** Detailed binding in the hexamer interface of CmTad1. Residues involved in hexamer formation are shown as sticks. Red dashed lines represent polar interactions. **f,** SLS studies of purified CmTad1 and its Q98A/E99A/M102A/W103A/K106A mutant. Calculated molecular weight is shown above the peaks. 5× mut represents the above mutant with 5 residues mutated. **g,** Overall structure of CmTad1 hexamer bound to cA_3_.Two views are shown. **h,** Detailed binding between CmTad1 and cA_3_. Residues involved in cA_3_ binding are shown as sticks. Red dashed lines represent polar interactions. 2Fo-Fc electron density of cA_3_ within one binding pocket is shown and contoured at 1 σ. **i,** Native PAGE showed the binding of CbTad1 and its mutants to cA_3_ and 1’’-2’ gcADPR. **j,** Overall structure of CmTad1 hexamer bound to cA_3_ and 1’’-3’ gcADPR. cA_3_ and 1’’-3’ gcADPR are shown as green and orange sticks, respectively. 2Fo-Fc electron density of cA_3_ and 1’’-3’ gcADPR within CbTad1 hexamer contoured at 1 σ.

### Tad1 forms a hexamer that binds cyclic trinucleotides and gcADPR with different binding sites

To understand how Tad1 interacts with CTNs, we first determined crystal structures of apo CmTad1 (2.56 Å) and its complex with cA_3_ (2.80 Å) and cAAG (3.27 Å), respectively (Extended Data Table 1). Surprisingly, in all the structures solved, CmTad1 is a hexamer (Figure 1c), rather than a dimer as previously proposed for CbTad1 ^4^. To verify the oligomeric state of Tad1 in solution, we performed static light scattering (SLS) analysis of CmTad1 and CbTad1, which also showed that both are hexamers in solution (Figure 1d). Then we re-examined the structural data of CbTad1 (PDB codes: 7UAV and 7UAW), which showed that a hexamer similar as that of CmTad1 could be generated by symmetry operations for both the apo CbTad1 (PDB code: 7UAV) and CbTad1 complexed with 1”-2’ gcADPR (PDB code: 7UAW) (Extended Data Figures 4a, b). The Tad1 hexamer can be viewed as a trimer of dimers with Dihedral D3 symmetry (Figure 1c). Analysis of the interface between two dimers showed that each protomer interacts with protomers from the other two dimers (Figure 1e). In CmTad1, R88 and K96 of protomer A interact with T95, Q98 and E99 of protomer C through polar interactions. In turn, Q98 and E99 of protomer A interact with R88 and K96 of protomer E. Meanwhile, M102, W103 and K106 of protomer A also interact with the same residues from protomer F through hydrophobic interactions (Figure 1e). Similar interactions can also be found in the CbTad1 hexamer (Extended Data Figure 4c). Notably, most of these interface residues are conserved among Tad1 homologs (Extended Data Figure 4d). Mutation of the interface residues of both CbTad1 and CmTad1 showed a significant backwards shift of the protein peak in gel filtration assay, and SLS analysis revealed that both mutant proteins are disrupted into dimers (Figure 1f, Extended Data Figure 4e). Structural superimposition showed that while both the N-terminal anti-parallel β-sheet and C-terminal two helices (α1 and α2) align well between CmTad1 and CbTad1, most of the loops linking the β-strands and helices display different conformations in the two proteins (Extended Data Figure 4f).

Next, we investigated the binding pockets of the CTNs. One Tad1 hexamer binds two CTNs with two distinct binding pockets that are far from the binding pockets for gcADPR molecules (Figure 1g). As expected, cA_3_ and cAAG bind at the same pocket in CmTad1 (Extended Data Figures 5a, 5b). In contrast to gcADPR molecules that bind in the pocket located in the interface of the dimer, CTNs are bound in the trimeric interface of the three Tad1 dimers (Figures 1g, Extended Data Figure 5c). Interestingly, the binding mode of CTNs in Tad1 is reminiscent of that of Acb2, which is also a hexamer and similarly binds two CTNs ^27^. As in Acb2, the CTN in Tad1 is also bound mainly through its three phosphate groups, each of which is coordinated by R88 of one CmTad1 protomer and T95 of another protomer through hydrogen bonds (Figure 1h). These two conserved residues are not only involved in binding of CTNs, but also hexamer formation of Tad1 (Extended Data Figure 4d). Consistently, mutation of either of the two corresponding residues in CbTad1, R90A and T97A, markedly reduced cA_3_ binding as revealed by native gel assay (Figure 1i). Moreover, C87, A91 and M92 from each CmTad1 protomer also form hydrophobic interactions with the bases of cA_3_. The binding mode of CTN indicates that hexamer formation is needed for its binding. As expected, CbTad1 and CmTad1 mutations that abolished the hexamer also lost the ability to bind CTNs (Extended Data Figure 6), supporting that hexamer formation is the prerequisite for cA_3_ binding.

To confirm that the binding sites of the CTNs and gcADPR molecules in Tad1 are independent of each other, we tested the binding of 1”-2’ gcADPR with the R90A or T97A CbTad1 mutant proteins. A native gel assay showed similar shifts of the two CbTad1 mutants as WT CbTad1 upon adding 1”-2’ gcADPR (Figure 1i), suggesting that binding to gcADPR is not affected by the two mutations. In turn, we tested whether binding of CTNs is affected by disruption of the binding sites of gcADPR. We also solved the structure of CbTad1 complexed with 1”-3’ gcADPR at 2.16 Å resolution (Extended Data Table 1), whose structure has been recently reported in a preprint, but not released by the Protein Data Bank ^5^. The structure showed that one CbTad1 hexamer binds six 1”-3’ gcADPR molecules (Extended Data Figure 7a) and the binding mode of 1”-3’ gcADPR is similar to that of 1”-2’ gcADPR (Extended Data Figure 7b) ^4^. Notably, mutation of the binding pocket R109A/R113A or F82A/N92A (Extended Data Figure 7b) exhibited a severely reduced binding to 1”-2’ gcADPR (Extended Data Figure 7c). However, these mutations did not interfere with the binding of cA_3_ (Extended Data Figure 7c). Taken together, these data collectively show that one Tad1 hexamer binds two CTNs through two pockets independent of those that bind gcADPR molecules.

Structural alignment between apo CmTad1 and its complexes with CTNs showed that the binding of CTNs does not induce a conformational change of CmTad1, with a root mean square deviation (RMSD) of 0.305 and 0.219 Å (Cα atoms) for CmTad1-cA_3_ and CmTad1-cAAG compared to the apo CmTad1, respectively (Extended Data Figure 5d, e). This suggests that Tad1 might be able to interact with CTNs and gcADPR molecules simultaneously. Therefore, we co-crystallized CbTad1 with both cA_3_ and 1”-3’ gcADPR and then solved its crystal structure at a resolution of 2.31 Å (Extended Data Table 1). The structure clearly showed that one CbTad1 hexamer binds to two cA_3_ and six 1”-3’ gcADPR molecules simultaneously (Figure 1j). Structural alignment between CbTad1-cA_3_-1”-3’ gcADPR and apo CbTad1 also showed little conformational changes except in the binding pocket of 1”-3’ gcADPR (Extended Data Figure 5f).

### Tad1 binds cyclic dinucleotides and gcADPR with the same binding pocket

To understand how CbTad1 interacts with CDNs, we determined the crystal structures of CbTad1 complexed with 2’,3’-cGAMP at 2.37 Å (Figure 2a and Extended Data Table 1). Surprisingly, 2’,3’-cGAMP binds in the same binding pocket as gcADPR molecules in CbTad1 (Figure 2b), and therefore a CbTad1 hexamer binds six 2’,3’-cGAMP molecules in total with two in each CbTad1 dimer (Figure 2a). Interestingly, comparison with the structure of CbTad1-1”-3’ gcADPR complex showed that in the binding pocket, the adenosine monophosphate moiety of 2’,3’-cGAMP almost completely overlaps with the corresponding part of 1”-3’ gcADPR (Figure 2b). On this side, the C-terminal residues 116-122 of CbTad1 also form an ordered lid to cover the 2’,3’-cGAMP molecule as in the CbTad1-1”-3’ gcADPR structure. However, the loop linking β4-α1 does not move to seal the binding pocket as it does in the CbTad1-1”-3’ gcADPR structure, instead keeps a similar conformation as the apo CbTad1 (Figure 2b). This makes sense because the same movement of this loop will cause steric clash to 2’,3’-cGAMP, especially that after the same movement R57 will even overlap with the guanine base of 2’,3’-cGAMP (Extended Data Figure 8a). In the binding pocket of 2’,3’-cGAMP, most of the residues that are involved in gcADPR binding also bind to 2’,3’-cGAMP, such as F82, N92, R109, and R113 (Figure 3c). Consistently, the R109A/R113A and F82A/N92A mutations of CbTad1 which disrupt 1”-3’ gcADPR binding (Extended Data Figure 7c) also severely reduced 2’,3’-cGAMP binding (Figure 2d). Moreover, on the side of the guanine base, Q8 forms a hydrogen bond with the oxygen atom of the base. The guanine base is also sandwiched by L13 of one protomer and L119 of the other protomer through hydrophobic interactions (Figure 2c).

**Figure 2.**
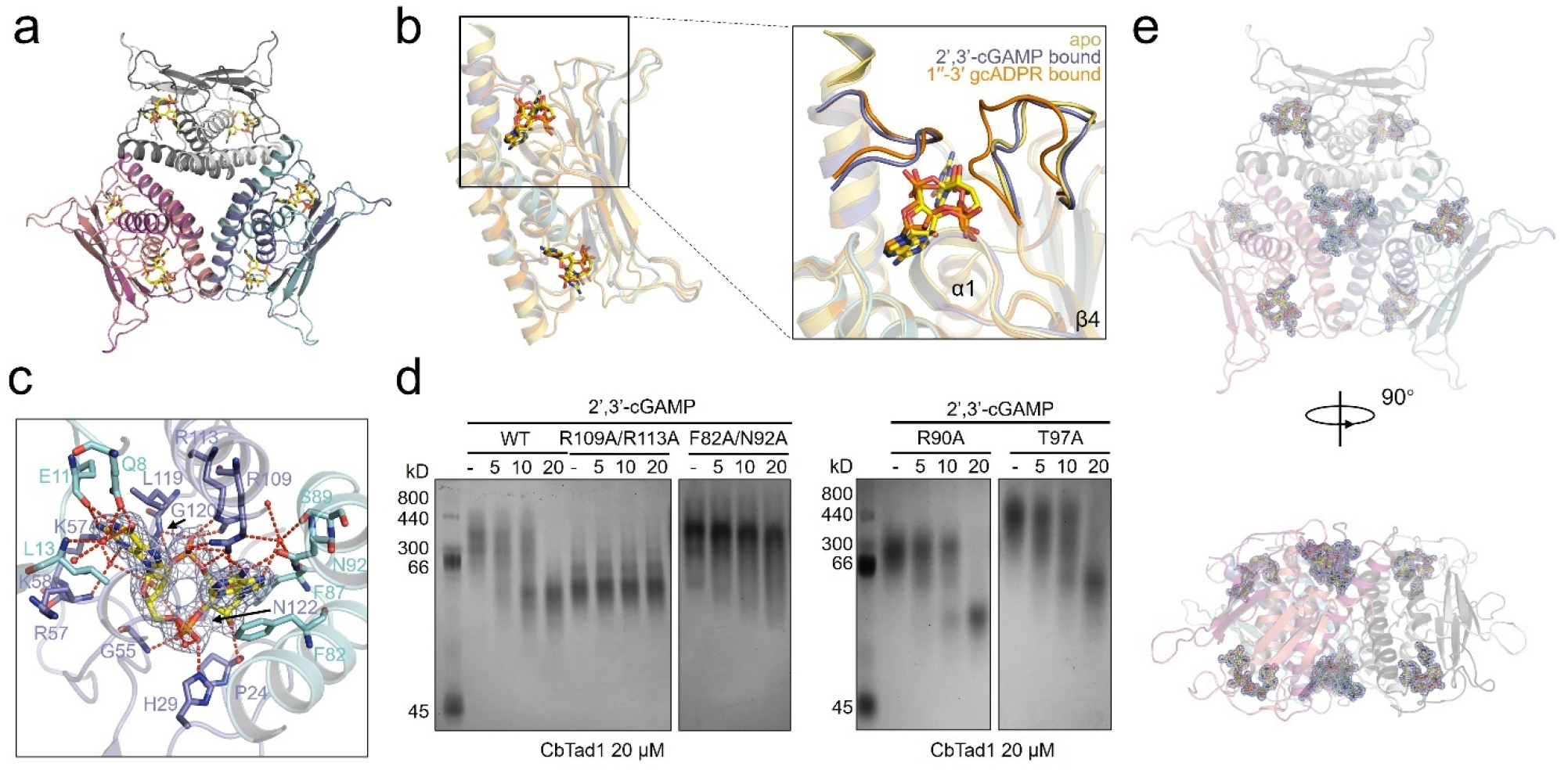
Tad1 binds to 2’,3’-/3’,2’-cGAMP using the same binding pocket as gcADPR molecules. **a**, Overall structure of CbTad1 hexamer bound to 2’,3’-cGAMP, which is shown as yellow sticks. **b,** Structural superimposition of apo, 1’’-3’ gcADPR-bound and 2’,3’-cGAMP-bound CbTad1 protein. 1’’-3’ gcADPR and 2’,3’-cGAMP are shown as orange and yellow sticks, respectively. The two loops that undergo conformational changes upon ligand binding are highlighted. **c,** Detailed binding between CbTad1 and 2’,3’-cGAMP. Residues involved in 2’,3’-cGAMP binding are shown as sticks. Red dashed lines represent polar interactions. 2Fo-Fc electron density of 2’,3’-cGAMP within one binding pocket is shown and contoured at 1 σ. **d,** Native PAGE showed the binding of CbTad1 and its mutants to 2’,3’-cGAMP. **e,** Overall structure of CmTad1 hexamer complexed with cA_3_ and 2’,3’-cGAMP. cA_3_ and 2’,3’-cGAMP are shown as green and yellow sticks, respectively. Two views are shown. 2Fo-Fc electron density of cA_3_ and 2’,3’-cGAMP within CbTad1 hexamer contoured at 1 σ.

**Figure 3.**
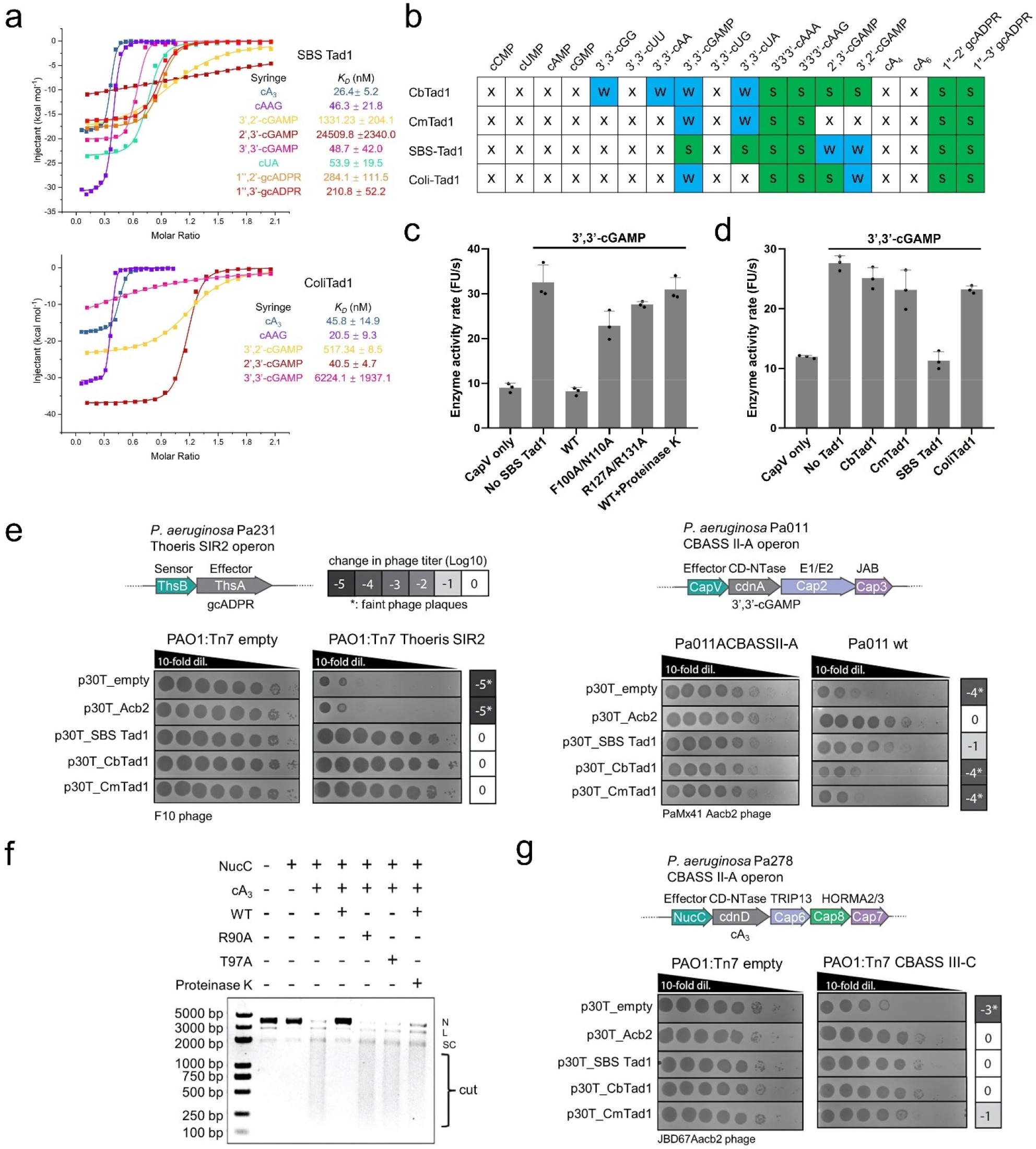
Tad1 antagonizes Type II-A and Type III-C CBASS immunity. **a**, ITC assays to test binding of cyclic oligonucleotides to SBS Tad1 and ColiTad1. Representative binding curves and binding affinities are shown. The *K_D_* values are mean ± s.d. (n=3). Raw data for these curves are shown in Extended Data Figure 2. **b,** Summary of the binding results of Tad1 homologs. Words in black: verified only by native PAGE. Χ: no binding; W: binding *K*_D_ higher than 400 nM. S: shift in native gel or binding *K*_D_ lower than 400 nM by ITC or SPR. **c,** CapV enzyme activity in the presence of 3’,3’-cGAMP and resorufin butyrate, which is a phospholipase substrate that emits fluorescence when hydrolyzed. The enzyme activity rate was measured by the accumulation rate of fluorescence units (FUs) per second. To test the effects of Tad1 homologs to sequester 3’,3’-cGAMP, Tad1 or its mutants (8 µM) was incubated with 3’,3’-cGAMP (0.8 µM) for 30 min. Filtered nucleotide products were used for the CapV activity assay. Data are mean ± SD (n=3). **d,** CapV enzyme activity with Tad1 homologs. The experiment was performed as in c. **e,** Plaque assays to test the activity of Tad1 against Thoeris and Type II-A CBASS immunity *in vivo*. Organization of *P. aeruginosa* Pa231 Thoeris and *P. aeruginosa* Pa011 CBASS II-A operons shown. F10 phage was spotted in 10-fold serial dilutions on a lawn of *P. aeruginosa* cells expressing Thoeris operon genes (PAO1:Tn7 Thoeris SIR2), or without Thoeris (PAO1:Tn7 empty). PaMx41Δ*acb2* was spotted on a lawn of Pa011 cells with deletion of CBASS operon (Pa011ΔCBASS II-A) or Pa011 wild type cells (Pa011 wt), electroporated with pHERD30T plasmids carrying Tad1 genes or empty vector. **f,** Effect of CbTad1 or its mutants on cA_3_-activated NucC effector protein function. After treatment with proteinase K, the released cA_3_ also showed the ability to activate the nuclease activity of NucC. The concentration of NucC, cA_3_, CbTad1 and proteinase K is 10 nM, 5 nM, 200 nM and 1 µM, respectively. N denotes nicked plasmid, SC denotes closed-circular supercoiled plasmid, and cut denotes fully digested DNA. **g,** Plaque assays to test the activity of Tad1 against Type III-C CBASS immunity *in vivo*. Organization of *P. aeruginosa* Pa278 Type III-C CBASS operon shown. JBD67Δ*acb2* phage was spotted in 10-fold serial dilutions on a lawn of *P. aeruginosa* cells expressing Pa278 CBASS operon genes (PAO1:Tn7 CBASS III-C), or without the system (PAO1:Tn7 empty), electroporated with pHERD30T plasmids carrying Tad1 genes or empty vector.

To understand why CbTad1 only shows high affinity to 2’,3’- and 3’,2’-cGAMP among the CDNs we tested, we performed docking studies of 3’,3’-/3’,2’-cGAMP into the binding pocket of 2’,3’-cGAMP in CbTad1 (Extended Data Figure 8b). Structurally, in 2’,3’- or 3’,2’-cGAMP, a 5’-GMP unit is connected with a 5’-AMP unit via a 2’-5’ (or 3’-5’) and a 3’-5’ (or 2’-5’) phosphodiester bond to form a cyclic structure, which result in the same structure of the cyclic phosphate-ribose backbone (Extended Data Figure 8b). However, in 3’,3’-cGAMP, a 5’-GMP unit is connected with a 5’-AMP unit with two 3’-5’ phosphodiester bonds, leading to a different structure of cyclic backbone, which might explain its low affinity to CbTad1. Moreover, cUA/cGG/cAA has the same cyclic backbone as 3’,3’-cGAMP and also has low affinity to CbTad1. Interestingly, while CbTad1 binds to multiple CDNs, CmTad1 only shows weak binding to 3’,3’-cGAMP and cUA. Comparison of the 2’,3’-cGAMP binding pocket between CbTad1 and CmTad1 showed that most of the loops surrounding 2’,3’-cGAMP display different lengths and conformations between the two proteins (Extended Data Figure 8c), and these loops of CbTad1 except the C-terminal loop do not move upon 2’,3’-cGAMP binding. Moreover, CbTad1 residues involved in binding to the 5’-GMP moiety of 2’,3’-cGAMP are also not conserved in CmTad1 (Extended Data Figure 4d). Similarly as gcADPR, 2’,3’-cGAMP binding does not induce a conformational change in the binding pocket of CTNs either (Extended Data Figure 8d). Moreover, mutation of the CTN binding residues does not affect binding of 2’,3’-cGAMP (Figure 2d) and vice versa (Extended Data Figure 7c). Then we moved on to co-crystallize CbTad1 with both cA_3_ and 2’,3’-cGAMP and then solved its crystal structure at a resolution of 1.54 Å (Extended Data Table 1), which clearly showed that a CbTad1 hexamer binds to two cA_3_ and six 2’,3’-cGAMP molecules simultaneously (Figure 2e). Structural alignment between CbTad1-cA_3_-2’,3’-cGAMP and apo CbTad1 also showed little conformational changes except in the binding pocket of 2’,3’-cGAMP (Extended Data Figure 8e).

### Cyclic dinucleotide binding spectra are different among Tad1 homologs

Different binding spectrum of CDNs between CbTad1 and CmTad1 indicates that Tad1 homologs might show different CDN binding spectra. Generation of a phylogenetic tree using PSI-BLAST to identify Tad1 homologs revealed numerous distinct clades of Tad1 and showed that CbTad1 and CmTad1 are represented on distant branches of the Tad1 phylogenetic tree (Extended Data Figure 9). Importantly, most Tad1 orthologs retain both CDN/gcADPR and CTN binding sites, whereas only 12% and 6% of proteins have substitutions in CDN/gcADPR or CTN binding sites, respectively (Extended Data Figure 9). To test the binding activities of diverse homologs, we purified Tad1 from *Bacillus cereus* phage SBSphiJ7 (named SBS Tad1) and *Colidexitribacter sp. OB.20* (named ColiTad1) and test their binding to the same array of cyclic oligonucleotides by native gel (Extended Data Figure 10) combined with ITC assays (Extended Data Figure 11). Both SBS Tad1 and ColiTad1 also bind to cA_3_/cAAG and cADPR isomers, demonstrating a broadly conserved function of this family. Interestingly, SBS Tad1 binds to 3’,3’-cGAMP/cUA with high affinity (*K*_D_ values of 48.7 and 53.9 nM, respectively) and 2’,3’-/3’,2’-cGAMP with low affinity (Figures 3a, Extended Data Figure 10, 11), which is the opposite of CbTad1. Notably, these *K*_D_ values are also comparable to the SBS Tad1 binding affinity for 1”-2’ and 1”-3’ gcADPR of 284 and 210 nM, respectively (Figures 3a, Extended Data Figure 11). Moreover, ColiTad1 binds to 2’,3’-cGAMP with high affinity and 3’,3’-/3’,2’-cGAMP with low affinity. Analysis of the well sequenced *Bacillus cereus* group, which is the bacterial hosts for SBS Tad1 revealed multiple commonly encoded CBASS CD-NTases (i.e. CdnB, CdnD, CdnE, and CdnG) that produce the spectrum of cyclic oligonucleotides that SBS Tad1 binds to (Extended Data Figure 3d). None of the known CD-NTases used in our search (see Methods) were identified in *Colidexitribacter* genomes. Taken together, these combined biochemical and bioinformatic results indicate that Tad1 homologs maintain a conserved ability to bind to CBASS CTN and gcADPR signals, with a variable spectrum of high affinity binding to CBASS CDNs (Figure 3b).

### Tad1 antagonizes Type II-A and Type III-C CBASS immunity

Since SBS Tad1 displays high affinity binding to 3’,3’-cGAMP, we tested whether SBS Tad1 can inhibit Type II-A CBASS immunity that uses 3’,3’-cGAMP signaling molecules to activate a phospholipase (CapV) effector protein. To this end, we first used *in vitro* CapV activity assay we set up in our previous study ^18^. While CapV activity could be activated by 3’,3’-cGAMP, its activity was abrogated when SBSTad1 was preincubated with 3’,3’-cGAMP (Figure 3c). Following proteolysis of SBS Tad1, the released molecule again activated the CapV activity. The SBS Tad1 mutants F100A/N110A and R127A/R131A, designed based on its conserved CDN binding sites, both displayed decreased 3’,3’-cGAMP binding (Extended Data Figure 12) and reduced inhibition on CapV activity (Figure 3c). Compared to SBS Tad1, other Tad1 homologs did not show significant inhibition of CapV activity, likely due to their weak binding to 3’,3’-cGAMP (Figure 3d). These results demonstrate that SBS Tad1 antagonizes Type II-A CBASS immunity *in vitro* through sequestering 3’,3’-cGAMP signaling molecules.

To determine whether SBS Tad1 can inhibit this same 3’,3’-cGAMP-based CBASS system *in vivo,* the different Tad1 proteins were expressed in the *Pseudomonas aeruginosa* strain BWHPSA011 (Pa011) with an active Type II-A CBASS system ^18^. We performed phage infection assays with the CBASS-targeted phage PaMx41 that lacks the anti-CBASS gene *acb2* (PaMx41Δ*acb2*). We observed that SBS Tad1, but not CmTad1 or CbTad1, which do not bind tightly to 3’,3’-cGAMP, inhibited Type II-A CBASS activity to nearly the same extent as the Acb2 positive control (Figure 3e). To confirm that all assayed proteins express well and retain anti-Thoeris activity *in vivo,* we identified a canonical Thoeris system in the *P. aeruginosa* strain MRSN390231 (Pa231) and expressed it from the chromosome of a strain that naturally lacks all known cyclic nucleotide signaling systems (PAO1). Expression of this Thoeris system reduced the titer of phage F10 by 5 orders of magnitude. However, co-expression of SBS Tad1, CmTad1, and CbTad1 inhibited Thoeris activity and rescued F10 phage titer, whereas Acb2 had no impact on Thoeris (Figure 3e). Acb2 also had no observed binding to the gcADPR molecules *in vitro* (Extended Data Figure 13). These data collectively demonstrates that the Pa231 Thoeris system uses a canonical signaling gcAPDR molecule and that Tad1 proteins antagonize canonical Thoeris, with one Tad1 homolog inhibiting a 3’3’-cGAMP CBASS system *in vivo*, consistent with *in vitro* binding patterns.

Since all Tad1 homologs tested display high affinity binding to CTNs, we tested whether Tad1 can inhibit Type III-C CBASS immunity that uses cA_3_ signaling molecules to activate a non-specific endonuclease (NucC) effector protein ^33,34^. Using the NucC enzyme from *P. aeruginosa* strain ATCC 27853 (Pa278), we showed that addition of cA_3_ activates the DNA cleavage activity, whereas adding WT CbTad1 significantly decreased NucC activity (Figure 3f). Moreover, following proteolysis of CbTad1, the released cA_3_ molecule again activated the NucC activity (Figure 3f). The R90A and T97A CbTad1 mutant proteins, which almost lost cA_3_ binding, displayed no inhibition of NucC activity. The same Pa278 Type III-C CBASS operon was chromosomally integrated into the PAO1 strain described above. CBASS Pa278 CBASS reduces the titer of phage JBD67Δ*acb2* by 3 orders of magnitude (Figure 3g). Co-expression of SBS Tad1, CbTad1, CmTad1, or an Acb2 control all fully inhibited cA_3_-based CBASS activity and rescued JBD67Δ*acb2* phage titer. Together, these data provide *in vivo* and *in vitro* evidence that Tad1 is a flexible anti-CBASS sponge protein, binding to both CDNs and CTNs involved in immunity, as well as an effective anti-Thoeris sponge protein.

### HgmTad2 sequesters multiple CBASS cyclic dinucleotides

Tad2 is a recently discovered anti-Thoeris sponge identified from *Bacillus cereus* phage SPO1 that works by sequestering gcADPR through a completely different structural fold from Tad1 ^5^. Using the same array of cyclic nucleotides that we used to study Tad1, we first tested SPO1 Tad2. While binding to gcADPR molecules was confirmed by native gel (Extended Data Figure 14a), no significant shift of SPO1 Tad2 was observed upon adding any of the other cyclic nucleotides (Extended Data Figure 14a), suggesting that SPO1 Tad2 might not bind any of these signaling molecules.

The Tad2 family of proteins is quite widespread in numerous MGEs (mobile genetic elements) and contains a domain of unknown function DUF2829. This domain was previously found in an anti-CRISPR (Acr), AcrIIA7, derived from <u>h</u>uman <u>g</u>ut <u>m</u>etagenomic libraries ^35^ (short for HgmTad2 hereafter). We decided to test whether HgmTad2 also sequesters gcADPR molecules. Interestingly, during purification, HgmTad2 eluted in three separate peaks in the process of ion exchange chromatography (Extended Data Figure 15a), which displayed different migrations in native gel. We collected the three components separately and tested whether they bind to gcADPR molecules. Native gel assay showed a shift of the purified HgmTad2 protein in all the three states upon adding 1’’-2’gcADPR (Extended Data Figure 15b), suggesting gcADPR binding. Then, we moved on to solve the structures of HgmTad2, HgmTad2-1’’-2’ gcADPR as well as HgmTad2-1’’-3’ gcADPR complexes. These complexes were obtained by expression of HgmTad2 alone or during co-expression with TIR protein from *Brachypodium distachyon*, or co-expression with ThsB from *Bacillus cereus* MSX-D12 ^4^, respectively (Extended Data Table 1). Purified HgmTad2 in the three different states were used separately during crystallization. Surprisingly, during structure solution, we found that a clear density with a shape similar to that of 3’,3’-cyclic di-GMP (cGG) was visible in all the solved structures using HgmTad2 of States 2 and 3 (Figure 4a), which simultaneously contained gcADPR molecules at a distinct site when co-expressed with gcADPR-producing enzymes. These enzymes and the significance of cGG as a CBASS signaling molecule will be discussed below. However, there was no such density in the structures using HgmTad2 in State 1. This suggested that HgmTad2 in States 2 and 3 contains cGG or other similar molecule bound during its expression in *E. coli*. To verify that the density corresponds to cGG, purified HgmTad2 in States 2 and 3 was denatured by heating and filtered to obtain the nucleotide within the protein. The filtered nucleotide showed a similar retention time as cGG, but markedly different from that of 3’,3’-cGAMP (Figure 4b), further supporting that the nucleotide within purified HgmTad2 in States 2 and 3 is cGG. However, the sample of HgmTad2 in State 1 after the same procedure showed no peak here (Figure 4b). Together, these findings show that HgmTad2 can bind cGG and gcADPR simultaneously, and a large fraction of the purified HgmTad2 contains bound cGG from expression in *E. coli*.

**Figure 4.**
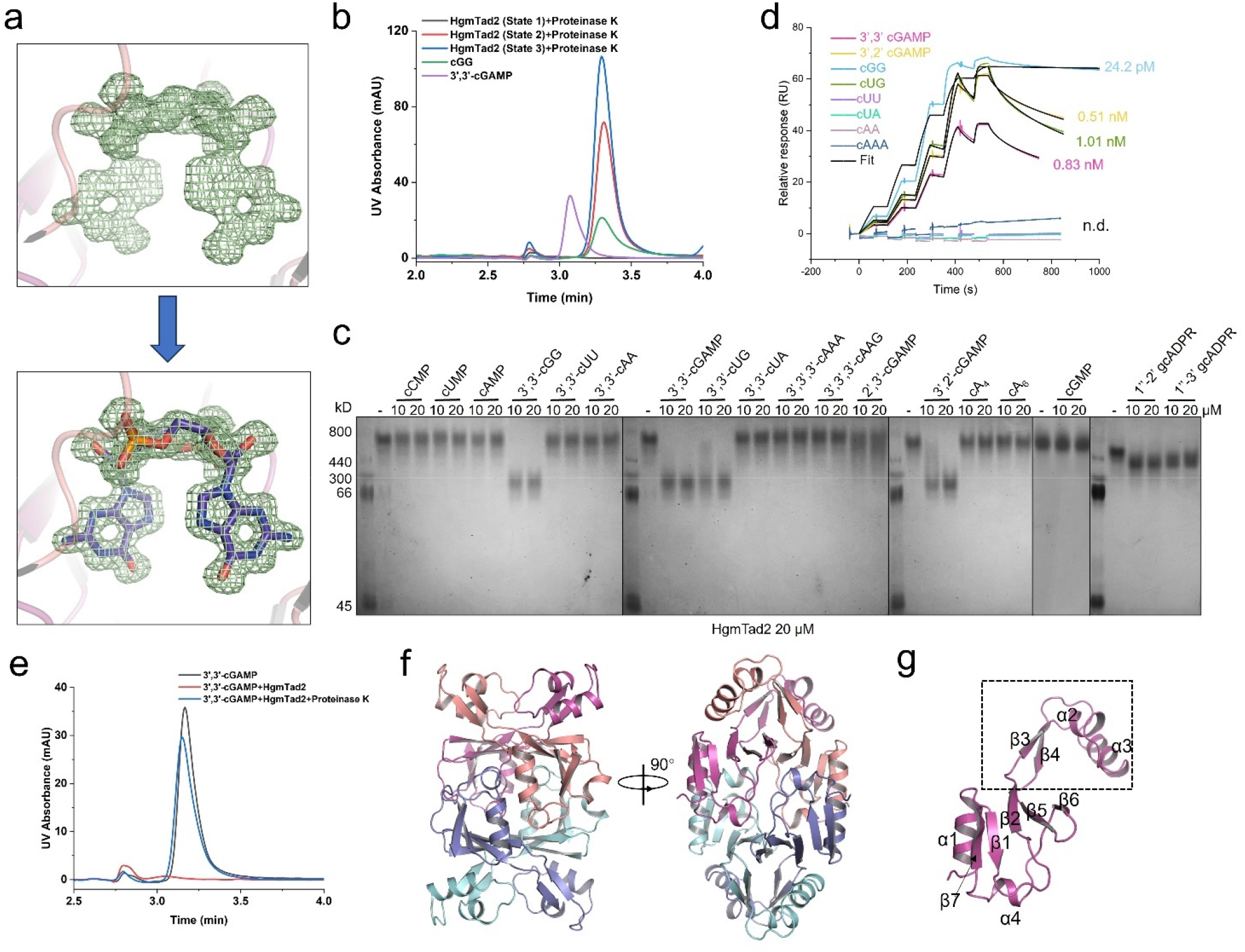
Tad2 binds an array of cyclic dinucleotides. **a**, The Fo-Fc density around the putative cGG in the structure of HgmTad2 of State 3 contoured at 2.5 σ. The density itself and with cGG placed are shown in the upper and lower panels, respectively. **b,** The molecules in HgmTad2 of three states released when treated with proteinase K was analyzed by HPLC. 3’,3’-cGAMP and cGG standard was used as controls. **c,** Native PAGE showed the binding of HgmTad2 of State 1 to cyclic oligonucleotides and gcADPR molecules. **d,** Overlay of sensorgrams from surface plasmon resonance (SPR) experiments, used to determine kinetics of HgmTad2 binding to CDNs. Data were fit with a model describing one-site binding for the ligands (black lines). **e,** The ability of HgmTad2 of State 1 to bind and release 3’,3’-cGAMP when treated with proteinase K was analyzed by HPLC. 3’,3’-cGAMP standard was used as a control. The remaining nucleotides after incubation with HgmTad2 was tested. **f,** Overall structure of HgmTad2 tetramer. Two views are shown. **g,** Structure of a protomer of HgmTad2. Secondary structures are labelled.

Binding of cGG by HgmTad2 was unexpected and this inspired us to consider that HgmTad2 might also bind other CDNs. To exclude the influence of bound cGG within HgmTad2 in binding assays, we only collected HgmTad2 in State 1 to test its binding spectrum using native gel assays, which showed a significant shift of HgmTad2 upon adding 3’,2’-/3’,3’-cGAMP/cGG/cUG and a minor shift upon adding 2’,3’-cGAMP, but no shift upon adding CTNs or other nucleotides (Figure 4c). These binding events were further verified by surface plasmon resonance (SPR) experiments, which showed that HgmTad2 binds to 3’,2’-/3’,3’-/2’,3’-cGAMP and cUG with a *K_D_* of ∼0.51, 0.83, 670 and 1.01 nM, respectively (Figures 4d, Extended Data Figure 15c). Surprisingly, the binding *K_D_* of cGG to HgmTad2 was calculated as high as 24.2 *p*M, possibly explaining why HgmTad2 stably bound to endogenous cGG during its expression in *E. coli*. HPLC assays demonstrated that HgmTad2 depletes 3’,3’-cGAMP, but doesn’t degrade it (Figure 4e), while SPO1 Tad2 does not deplete the molecule (Extended Data Figure 11b, c). Taken together, these results demonstrate that HgmTad2 specifically sequesters multiple CDNs used in CBASS immunity in addition to gcADPR molecules.

### Tad2 binds cyclic dinucleotides and gcADPR with different binding pockets

Next, we will introduce the structures of HgmTad2 and its complexes with gcADPR and CDNs. We solved in total six structures of HgmTad2, which are in apo form (1.70 Å), 1’’-2’ gcADPR-bound (2.10 Å), cGG-bound (1.38 Å), 3’,3’-cGAMP-bound form (2.11 Å), 1’’-2’ gcADPR-cGG-bound (2.28 Å) and 1’’-3’ gcADPR-cGG-bound (1.98 Å) forms, respectively. Since SPO1 Tad2 structure has not been released by the Protein Data Bank ^5^, we also solved its structure (2.27 Å) to compare with HgmTad2. HgmTad2 forms a tetramer similarly to SPO1 Tad2 (Figure 4f), and the tetrameric state of HgmTad2 and SPO1 Tad2 were also verified by SLS analysis (Extended Data Figure 16a). A Tad2 tetramer can be viewed as a dimer of dimers. Two Tad2 protomers interlock with each other to form an “X”-shaped dimer with a buried surface of ∼1400 Å^2^. And then, two such dimers further interlock with each other along the axis of helix α1 to form a tetramer, in which each protomer of the dimeric unit interacts with the two protomers within the other unit (Figure 4f). Each HgmTad2 protomer also contains an N-terminal α helix (α1) followed by an antiparallel five-stranded β sheet (β1–2, β5-7) as SPO1 Tad2 (Figure 4g). However, in the loop region linking β2 and β5, there are two α helices (α2-α3) and two β strands (β3-β4) in HgmTad2 (Figure 4g, Extended Data Figure 16b), compared to only one α helix in the corresponding loop region of SPO1 Tad2 ^5^. The C-terminal α helix (α4) of HgmTad2 is located between β6-β7.

Both SPO1 Tad2 and HgmTad2 tetramer bind two gcADPR molecules with two identical binding pockets, which are located in the middle region of the tetramer at the interface of two protomers from different dimeric units (Figure 5a). The gcADPR ligands are surrounded by loop L12 (between β1– β2), helix α4 and the loop linking β6 and α4 of one protomer (a), and loop L12, L56, α4 and β6 of the other (b) (Extended Data Figure 17a). Interestingly, the region between β1–β2 has two different conformations in the apo HgmTad2 structure: Two protomers that will together bind one gcADPR both form an extra helix (residues 19-24) away from each other in this region. The other two protomers both form a loop much nearer to each other (Extended Data Figure 17b). Interestingly, in the gcADPR-bound structure, all the four protomers form a loop in this region similar to that in the apo form (Extended Data Figure 17b), suggesting that this region of HgmTad2 is flexible and binding of gcADPR ligands will induce and stabilize it as a loop covering the ligand. Specifically, for 1’’-2’ gcADPR, its adenine base is coordinated by hydrogen bonds from T92_a_ and water-mediated hydrogen bonds from T82_b_, N22_a_ and mainchain oxygen and nitrogen atoms of L88_b_. Moreover, the adenine base is also stabilized by hydrophobic interactions from M76_a_, A78_a_ and V84_a_ (Figure 5b). The diphosphate backbone is bound by N22 and G23 from both protomers. The free hydroxyls in the ribose–ribose linkage are coordinated by hydrogen bonds from W87_a_, L88_a_, D93_a_ and D93_b_ (Figure 5b). Supporting this, mutations W21A/N22A and S90A/T92A/D93A of HgmTad2 markedly reduced both its binding to 1’’-2’ gcADPR (Extended Data Figure 17c) and its inhibition effect on 1’’-2’ gcADPR-activated NADase activity of ThsA (Figure 5c).

**Figure 5.**
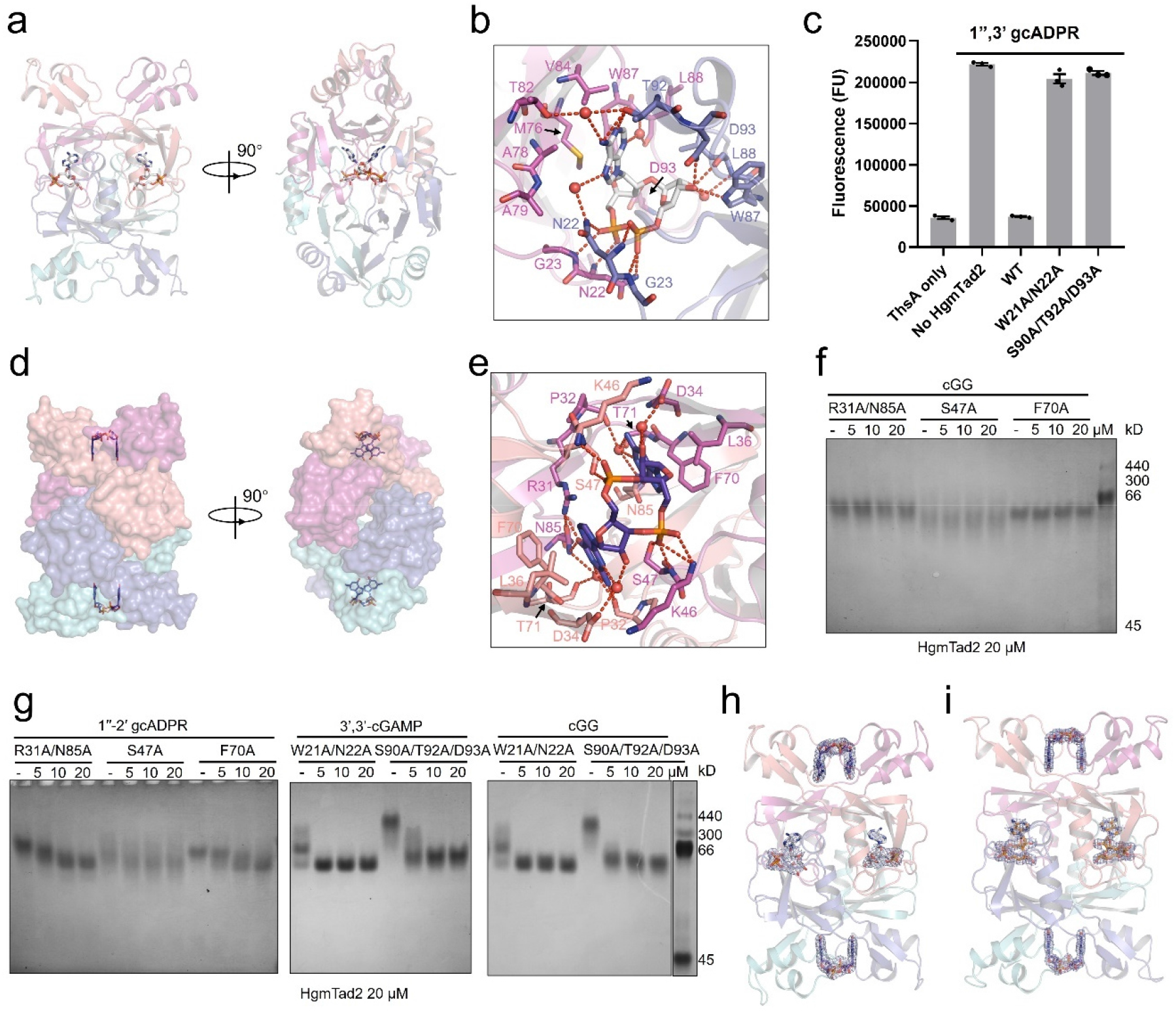
Tad2 binds cyclic dinucleotides and gcADPR molecules simultaneously. **a**, Overall structure of HgmTad2 tetramer bound to 1’’-2’ gcADPR, which is shown as gray sticks. **b,** Detailed binding between HgmTad2 and 1’’-2’ gcADPR. Residues involved in ligand binding are shown as sticks. Red dashed lines represent polar interactions. **c,** ThsA enzyme activity in the presence of 1’’-3’ gcADPR and ε-NAD. Wild-type (WT) and mutated HgmTad2 at 40 nM were incubated with 5 nM 1’’-3’ gcADPR. And then the reactions were filtered and their ability to activate ThsA NADase activity was measured. Bars represent the mean of three experiments, with individual data points shown. Data are mean ± SD (n=3). **d,** Overall structure of HgmTad2 tetramer bound to cGG, which is shown as purple sticks. HgmTad2 is shown as surface model. **e,** Detailed binding between HgmTad2 and cGG. Residues involved in ligand binding are shown as sticks. Red dashed lines represent polar interactions. **f,** Native PAGE showed the binding of HgmTad2 mutants to cGG. **g,** Native PAGE showed the binding of HgmTad2 mutants to 1’’-2’ gcADPR, 3’,3’-cGAMP or cGG. **h-i,** Overall structure of HgmTad2 tetramer bound to cGG and 1’’-2’ gcADPR simultaneously (**h**), or cGG and 1’’-3’ gcADPR simultaneously (**i**), cGG, 1’’-2’ gcADPR and 1’’-3’ gcADPR are shown as purple, gray and orange sticks, respectively. 2Fo-Fc electron density of the ligands within HgmTad2 tetramer is contoured at 1 σ.

HgmTad2 additionally binds to CDNs 3’,3’-cGAMP/cGG/cUG and 3’,2’-cGAMP in a distinct region of the protein (Figure 5d). An HgmTad2 tetramer binds two CDNs with two identical binding pockets, which are located at the top and bottom ends of the tetramer at the interface of two protomers within one dimeric unit (Figure 5d). As expected, cGG and 3’,3’-cGAMP bind at the same binding pocket (Extended Data Figure 17d, e). Each pocket is a symmetrical one, surrounded by the loop between α2 and α3, β2, β3 and the loop between them, as well as the C-terminal residue of β6 from both protomers (Extended Data Figure 17f). Notably, almost all of these are structural elements in the insertion (residues 32-72) between β2 and β5 of HgmTad2 (Extended Data Figure 16b), which is nearly twice the size of that in SPO1 Tad2 (residues 36-59). Binding of cGG causes some conformational changes to the structural elements surrounding the molecule in the binding pocket (Extended Data Figure 17g). In the HgmTad2-cGG complex, ligand binding is mediated by extensive hydrophobic and polar interactions. The guanine base is stabilized by hydrophobic interactions from L36 and F70. Moreover, it is coordinated by hydrogen bonds from R31 and N85 from one protomer, and mainchain carbonyls of P32 from the other protomer (Figure 5e), as well as water-mediated interactions from D34 and main chain carbonyls of T71 from the other protomer (Figure 5e). However, for the adenine base of 3’,3’-cGAMP, only one water mediated interaction can be formed by HgmTad2 apart from hydrophobic interactions from L36 and F70 (Extended Data Figure 17h), which may explain the high binding affinity of cGG and the inability of binding to cAA by HgmTad2. For the phosphate-ribose backbone, the phosphate group is coordinated by polar interactions from S47 and mainchain nitrogen atom of K46. To verify these residues, we mutated interacting residues of HgmTad2 (S47A, F70A, R31A/N85A) and tested their binding to both cGG and 3’,3’-cGAMP. Consistently, native gel showed no shift of these mutants upon adding either cGG or 3’,3’-cGAMP (Figure 5f, Extended Data Figure 17i).

To further confirm that the binding sites of the cyclic dinucleotides and gcADPR in HgmTad2 are independent of each other, we tested the binding of 1”-2’ gcADPR with S47A, F70A and R31A/N85A mutant proteins, as well as the binding of cGG/3’,3’-cGAMP with W21A/N22A and S90A/T92A/D93A mutant proteins. The results showed that mutation of either binding site does not decrease the binding of the other ligand (Figure 5g). This is also consistent with the fact that we obtained the co-structures of HgmTad2-1’’-2’-gcADPR-cGG (Figure 5h) and HgmTad2-1’’-3’-gcADPR-cGG (Figure 5i). Taken together, an HgmTad2 tetramer can bind to two cyclic dinucleotides and two gcADPR molecules simultaneously.

### Tad2 binds cyclic dinucleotides with its insertion domain

As mentioned above, HgmTad2 binds CDNs with its insertion between β2 and β5, which region is much shorter in SPO1 Tad2. Structural superimposition shows that while HgmTad2 and SPO1 Tad2 are similar in the gcADPR-binding domain, they are highly different in the CDN binding domain (Figure 6a). The insertion domain of HgmTad2 stretches out through an anti-parallel β sheet (β3-4) to create a cavity for binding of cyclic dinucleotides. However, the insertion in SPO1 Tad2 is much smaller and displays a highly different conformation (Figure 6a). Since the binding to cGG results from the insertion between β2 and β5 of HgmTad2 (Extended Data Figure 16b), we performed a sequence-based analysis to search for Tad2 homologs with long insertions like HgmTad2 that may enable binding to CDNs (Figure 6b, Extended Data Figure 18). Interestingly, two of such Tad2 homologs from *Sphingobacterium thalpophilum* (SptTad2) and *Salegentibacter* sp.BDJ18 (SaTad2) with similarly long insertions also bind to CDNs with different affinities (Figures 6c-e, Extended Data Figure 19). Notably, SptTad2 also binds cGG with a high affinity of 0.23 nM (Figure 6c), and purified SptTad2 also contains cGG bound during expression. To investigate whether SptTad2 uses a similar binding mode to bind CDNs, we solved the structure of SptTad2 bound with cGG (Figure 6f). Structural superimposition showed that the cGG binding pocket is highly similar between HgmTad2 and SptTad2 (Figure 6g), thereby demonstrating that Tad2 homologs can bind CDNs with their large insertion domain (35-41 residues) between β2 and β5. Interestingly, phylogenetic analysis showed that the organization of this domain is highly variable in distant Tad2 homologs, which might reflect their different CDN binding activity. The gcADPR binding site, however, is highly conserved with only 1% of Tad2 proteins predicted to be non-functional in gcADPR binding (only short Tad2 versions of 120-170 amino acids length were used for analysis) (Extended Data Figure 18).

**Figure 6.**
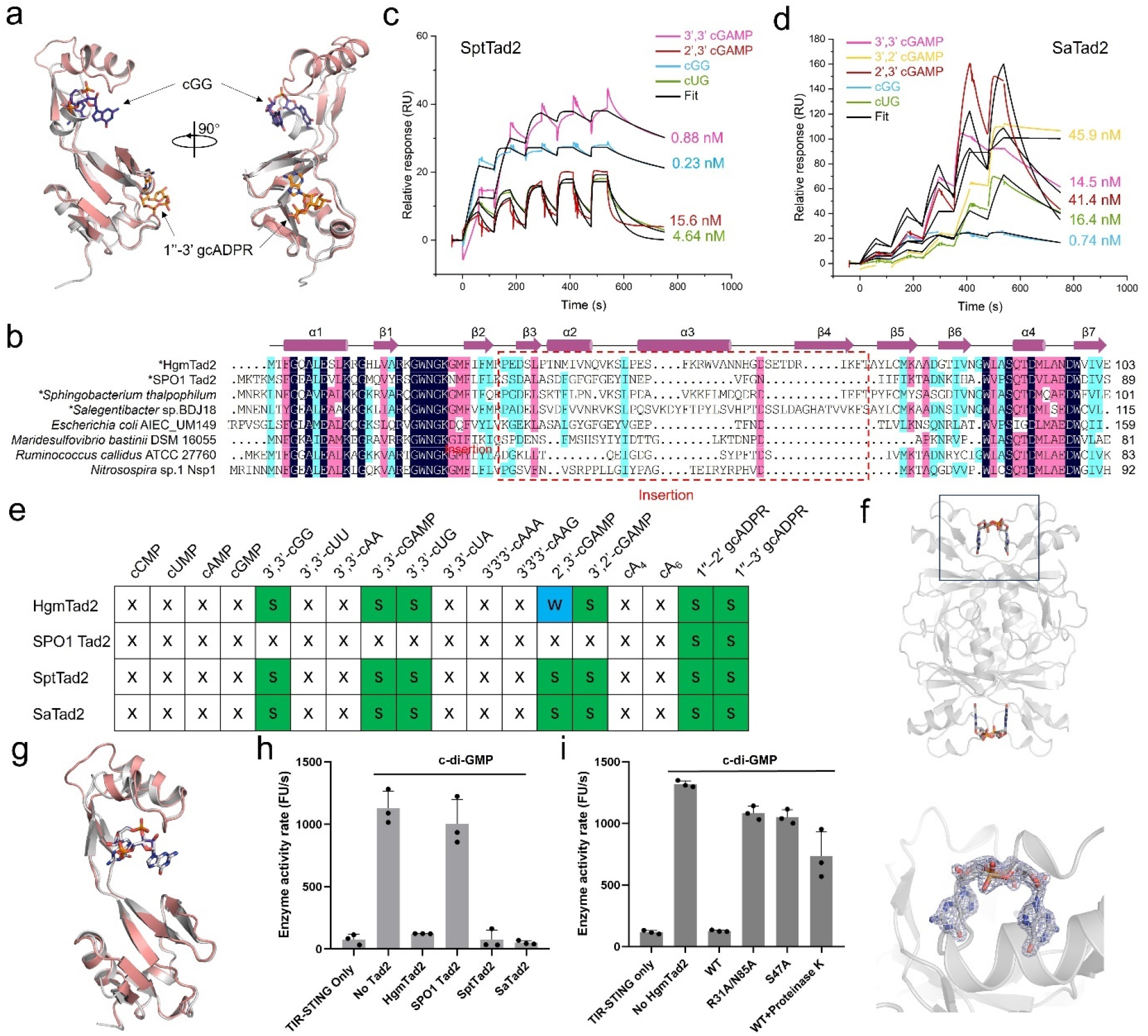
Tad2 antagonizes Type I-D CBASS immunity that uses cGG. **a**, Structural superimposition between HgmTad2-cGG-1’’-3’-gcADPR and SPO1 Tad2. HgmTad2 and small molecules are colored as in Fig. 6I. SPO1 Tad2 is colored gray. **b,** Sequence alignment among Tad2 homologs. Residues with 100 % identity, over 75 % identity and over 50 % identity are shaded in dark blue, pink and cyan, respectively. Secondary structural elements of HgmTad2 are shown above the alignment. The insertion region (residues 32-72) between β2 and β5 of HgmTad2 or between β2 and β3 of SPO1 Tad2 (residues 36-59) is marked with a rectangle. Biochemically studied Tad2 homologs are marked with an asterisk before its species name. **c-d,** SPR assay of SptTad2 (**c**) and SaTad2 (**d**). **e,** Summary of the binding results of Tad1 homologs. The figure is labelled as in Figure 3b. **f,** Overall structure of SptTad2 bound to cGG. A close view of the bound cGG with 2Fo-Fc electron density contoured at 1 σ is shown in the lower panel. **g,** Structural superimposition between HgmTad2-cGG and SptTad2-cGG. HgmTad2 and cGG are colored as in Fig. 6I. is colored gray. SptTad2 and its bound cGG are colored gray. **h, i,** TIR-STING NAD^+^ cleavage activity in the presence of cGG and nicotinamide 1,N^6^-ethenoadenine dinucleotide (εNAD), which emits fluorescence when cleavage. The enzyme activity rate was measured by the accumulation rate of fluorescence units (FUs) per second. To test the effects of HgmTad2 or its homologs to bind cGG, HgmTad2 or its homologs (1 μM) was incubated with cGG (50 nM) for 20 min. To test the effects of HgmTad2 or its mutants to bind and release cGG, HgmTad2 or its mutants (200 nM) was incubated with cGG (50 nM) for 20 min and then proteinase K (28.3 μg/mL) was added to release the nucleotide from the HgmTad2 protein, Filtered nucleotide products were used for the TIR NADase activity assay. Data are mean ± SD (n=3).

### Tad2 antagonizes Type I-D and Type II-A CBASS *in vitro*

Diguanylate cyclases (DGCs) produce cGG in many Gram-negative bacteria that are distinct enzymes from the CD-NTases that make CBASS nucleotides. cGG signaling in many bacteria controls motility and biofilm formation ^36^. Interestingly, HgmTad2 is encoded by *Bacteroides* phages. In *Bacteroides*, these EAL-containing DGCs are absent, but instead cGG signaling is mediated by a CBASS CD-NTase (CdnE) that signals to a TIR- or TM-STING fusion effector protein ^37^. Upon further investigation of *Bacteroides* and *Sphingobacterium* genomes, which are the bacterial hosts of HgmTad2- and SptTad2-encoding phage, respectively, we found that the bacteria both encode CBASS CD-NTases that are known to produce cGG (CdnE and CdnB) (Extended Data Figure 3b, c) ^21,37^, providing a biologically necessary role for these Tad2 proteins to strongly bind and sequester cGG signaling molecules. As such, we tested whether HgmTad2 or SptTad2 can inhibit Type I-D CBASS immunity that uses cGG signaling molecules with a previously reported TIR-STING activity assay ^37^. While activity of TIR-STING from *Sphingobacterium faecium* DSM 11690 could be activated by cGG, its activity was abrogated when HgmTad2 or SptTad2 was preincubated with cGG (Figure 6h). The R31A/N85A, S47A, and F70A HgmTad2 mutant proteins, which exhibited decreased cGG binding, displayed reduced inhibition of TIR-STING activity (Figure 6g). Moreover, following proteolysis of HgmTad2, the released cGG again partially activated the TIR-STING activity (Figure 6i). These results demonstrate that Tad2 antagonizes Type I-D CBASS immunity *in vitro* through sequestering cGG molecules.

Since HgmTad2/SptTad2/SaTad2 also display high affinity binding to 3’,3’-cGAMP, we tested whether they inhibit Type II-A CBASS immunity using the previous mentioned CapV activity assay. While CapV activity is activated by 3’,3’-cGAMP, it was abrogated when the cGG-free form of HgmTad2 or SptTad2 or SaTad2 was preincubated with 3’,3’-cGAMP (Extended Data Figure 20a). The R31A/N85A, S47A and F70A HgmTad2 mutant proteins, which exhibited decreased 3’,3’-cGAMP binding, also displayed reduced inhibition of CapV activity (Extended Data Figure 20b). Moreover, following proteolysis of HgmTad2, the released 3’,3’-cGAMP molecule again activated the CapV activity (Extended Data Figure 20b). These results demonstrate that Tad2 antagonizes Type II-A CBASS immunity *in vitro* through sequestering the 3’,3’-cGAMP molecule. Despite the Tad2 proteins inhibiting Thoeris activity *in vivo* (Extended Data Figure 20c), we did not observe inhibition of the Pa011 Type II-A CBASS activity *in vivo*, which is likely because the 3’,3’-cGAMP binding site is saturated with the highly abundant and common cGG nucleotide in *P. aeruginosa* (Extended Data Figure 20d).

### Tad2 does not antagonize SpyCas9 activity

We have demonstrated that HgmTad2 could simultaneously inhibit CBASS and Thoeris immunity as a sponge protein with two different binding pockets. However, this protein has been previously identified as an anti-CRISPR (Acr) protein, AcrIIA7 ^35^, whose inhibitory mechanism is unknown. HgmTad2 was previously shown to not interact with SpyCas9, but somehow inhibit its activity. It seemed surprising to us that this protein might have three inhibitory activities. Therefore, to query this activity and investigate the anti-CRISPR mechanism of HgmTad2, we first repeated the *in vitro* SpyCas9 cleavage assay in Uribe et al. 2019. Despite many trials and optimization of the reaction system, we still did not see Acr activity of HgmTad2 or the other Tad2 homologs in this study where AcrIIA11 successfully inhibits SpyCas9-mediated DNA cleavage (Extended Data Figure 20e). Consistent with this, chromosomal integration of SpyCas9 into the *P. aeruginosa* strain PAO1 demonstrated that the JBD30 phage is targeted ^38^, but HgmTad2 did not exhibit Acr activity (Extended Data Figure 20f). Taken together, our data suggests that HgmTad2 inhibits CBASS and Thoeris, but not the CRISPR-Cas9 system.

## Discussion

Thoeris and CBASS are two anti-phage systems that use different signaling molecules to mediate immunity. Two anti-immune proteins, Tad1 and Tad2, have been identified for Thoeris system and two anti-immune proteins, Acb1 and Acb2, for CBASS. Here, we demonstrated that anti-Thoeris proteins Tad1 and Tad2 also inhibit CBASS systems, which are generally more common, by sequestering a broad array of CDNs and CTNs. Therefore, Tad1 and Tad2 are the first phage-encoded sponge proteins that sequester multiple signaling molecules that are involved in two different anti-phage immune systems. Notably, Tad1 and Tad2 sequester cyclic oligonucleotides with completely distinct mechanisms. Tad1 is a hexamer that is assembled as a trimer of dimers. One Tad1 hexamer sequesters two CTNs using two separate pockets formed only in the case of the hexameric assembly, in which each pocket is composed of three interlocking protomers. In addition to CTNs, Tad1 also sequesters CDNs using the same binding pocket as gcADPR molecules. By contrast, Tad2 is a tetramer that binds two CDNs and two gcADPR molecules simultaneously. The binding pocket of CDNs in Tad2 is far from that of gcADPR and is also different from those of other known CDN binding proteins. Among the CDNs tested, HgmTad2 binds strongly to 3’,3’-cGAMP/cGG/cUG and 3’,2’-cGAMP, and weakly to 2’,3’-cGAMP. Notably, both Tad1 and Tad2 sequester 3’,2’-cGAMP, a signaling molecule that is not cleaved by Acb1, but has been recently implicated in both CBASS and cGAS-like signaling systems in eukaryotes ^22,39,40^.

Surprisingly, HgmTad2 displayed a *p*M-range binding affinity to cGG, which is much higher than any reported binding affinities to CDNs, and also much higher than affinities of HgmTad2 to other CDNs (Figure 4d). This explains why purified HgmTad2 contains cGG that is bound during its expression in *E. coli*. Moreover, cGG is a molecule that is not cleaved by Acb1 nor sequestered by Acb2. Therefore, to our knowledge, HgmTad2 is the first phage anti-immune protein to act as a cGG sponge, which might provide a useful reagent for studying cGG signaling not related to phage defense. In bacteria, cGG is the most widespread CDN that functions as a signaling molecule, regulating multiple aspects of bacterial growth and behavior, including motility, virulence, biofilm formation, and cell cycle progression ^36^. In *Bacteroides,* however, there are at least two known CD-NTases that produce cGG as a signaling molecule in CBASS immunity ^21,37^. Bioinformatic analyses demonstrate that cGG-based CBASS immunity is found in bacteria that HgmTad2 and SptTad2-encoding phages may infect.

While the binding mode of CTNs is similar between Tad1 and Acb2, the assembly mechanism of the hexamer and residues involved in binding are different between these proteins ^27^. Furthermore, our recent study on Acb2 demonstrates that it functions as a sponge that binds to both CDNs and CTNs used by a single bacterial anti-phage system. However, in the present study, we report that both Tad1 and Tad2 are sponge proteins that bind to a broad array of cyclic oligonucleotides from two independent anti-phage systems. Therefore, since bacterial species may contain both CBASS and Thoeris systems ^11,29^, Tad1 and Tad2 represent a unique class of proteins that are advantageous over the pan-immune arsenal of their host. Altogether, our findings demonstrate the remarkable potency of two anti-immune sponge proteins. Together with Acb2, these new data on Tad1 and Tad2 establish a paradigm of anti-immune sponge proteins with >1 binding site. We predict that a broad distribution of anti-immune sponges with multiple binding sites for signaling molecules may exist for anti-viral immune signaling systems across all domains of life.

## Acknowledgments

We thank the staff at beamlines BL02U1 and BL19U1 of the Shanghai Synchrotron Radiation Facility for their assistance with data collection. We thank the Tsinghua University Branch of China National Center for Protein Sciences Beijing and Dr. Shilong Fan for providing facility support for X-ray diffraction of the crystal samples. We thank Drs. Yuanyuan Chen, Zhenwei Yang, Bingxue Zhou at the Institute of Biophysics, Chinese Academy of Sciences for technical help with ITC and SPR experiments. Y. F. is supported by National key research and development program of China (2022YFC3401500 and 2022YFC2104800), the National Natural Science Foundation of China (32371329 and 32171274), Beijing Nova Program (20220484160) and the Fundamental Research Funds for the Central Universities (QNTD2023-01). E.H. is supported by the National Science Foundation Graduate Research Fellowship Program [Grant No. 2038436]. Any opinions, findings, and conclusions or recommendations expressed in this material are those of the authors and do not necessarily reflect the views of the National Science Foundation. J.B.-D. is supported by the National Institutes of Health [R21AI168811, R01GM127489], the Vallee Foundation, and the Searle Scholarship.

## Author contributions

Y.F. and J.B.-D. conceived and supervised the project and designed experiments. D.L., W.X., Y.W., X.L., Z.G., X.Z. and X.C. purified the proteins, grew and optimized the crystals, collected the diffraction data and performed *in vitro* activity analysis and binding assays. Y.X. solved the crystal structures with the help of Y.F. and Y.Z.. I.F. performed *in vivo* phage experiments, strain engineering, and Tad1/Tad2 protein bioinformatics. J. R. performed HPLC assays. E.H. executed cyclase bioinformatics. Y.F. wrote the original manuscript. J.B.-D., Y.F., I.F., and E.H. revised the manuscript.

## Declaration of interests

J.B.-D. is a scientific advisory board member of SNIPR Biome and Excision Biotherapeutics, a consultant to LeapFrog Bio and BiomX, and a scientific advisory board member and co-founder of Acrigen Biosciences. The Bondy-Denomy lab received research support from Felix Biotechnology.

## Data Availability

The accession numbers for the coordinate and structure factors reported in this paper are PDB: 8KBB (apo-CmTad1), 8KBC (CmTad1-cA_3_), 8KBD (CmTad1-cAAG), 8KBE (CbTad1-1’’,3’-gcADPR), 8KBF (CbTad1-1’’,3’-gcADPR-cA_3_), 8KBG (CbTad1-2’,3’-cGAMP), 8KBH (CbTad1-2’,3’-cGAMP-cA_3_), 8KBI (apo-HgmTad2), 8KBJ (HgmTad2-1’’,2’-gcADPR) 8KBK (HgmTad2-1’’,2’-gcADPR-cGG), 8KBL (HgmTad2-1′′-3′-gcADPR-cGG), 8KBM (HgmTad2-cGG), 8WJC (HgmTad2-3’,3’-cGAMP), 8WJD (SptTad2-cGG) and 8WJE (apo-SPO1 Tad2). This paper does not report original code. Any additional information required to reanalyze the data reported in this paper is available from the corresponding authors upon request.

## Materials and Methods

### Bacterial strains and phages

The *P. aeruginosa* strains (BWHPSA011, ATCC 27853, MRSN390231, PAO1) and *E. coli* strains (DH5ɑ) were grown in Lysogeny broth (LB) medium at 37°C both with aeration at 225 r.p.m. Bacteria plating was performed on LB broth supplemented with gentamicin for maintaining pHERD30T plasmid (50 µg ml^-1^ for *P. aeruginosa* and 20 µg ml^-1^ for *E. coli)*, as well as with 10 mM MgSO_4_ for phage spot assays. Gene expression in *P. aeruginosa* was induced by the addition of 0.2% L-arabinose or 0.3 mM isopropyl-β-D-thiogalactopyranoside IPTG unless stated otherwise. The *E. coli* BL21 (DE3) strain was used for recombinant protein overexpression and grown in Lysogeny broth (LB) medium. The cells were grown at 37°C until OD_600nm_ reached 0.8 and then induced at 18°C for 12 h.

### Protein expression and purification

The *Clostridium botulinum* Tad1, *Clostridioides mangenotii* Tad1, *SBSphiJ7* Tad1, *Colidexitribacter* Tad1, *Sphingobacterium thalpophilum* Tad2, *Salegentibacter* sp. BDJ18 Tad2, SPO1 Tad2, *P. aeruginosa* BWHPSA011 CapV, *P. aeruginosa* ATCC 27853 NucC, *Bacillus cereus* MSX-D12 ThsA, *Sphingobacterium faecium* DSM 11690 STING and *S. pyogenes* Cas9 genes were synthesized by GenScript and codon-optimized for expression in *E. coli*. The full-length CmTad1, SBS Tad1, ColiTad1, SptTad2, SaTad2, EcTad2, ThsA, CapV, NucC, *Sf*STING and SpyCas9 gene was amplified by PCR and cloned into a modified pET28a vector in which the expressed protein contains a His6 tag or His6-SUMO tag. The full-length CbTad1 gene was amplified by PCR and cloned into a modified pRSFDuet vector in which the expressed CbTad1 protein contains a His6 tag. The Tad1 or Tad2 mutants were generated by two-step PCR and were subcloned, overexpressed and purified in the same way as wild-type protein. All the proteins were expressed in *E. coli* strain BL21 (DE3) and induced by 0.2 mM isopropyl-β-D-thiogalactopyranoside (IPTG) when the cell density reached an OD_600nm_ of 0.8. After growth at 18°C for 12 h, the cells were harvested, resuspended in lysis buffer (50 mM Tris–HCl pH 8.0, 300 mM NaCl, 10 mM imidazole and 1 mM PMSF) and lysed by sonication. The cell lysate was centrifuged at 20,000 g for 50 min at 4°C to remove cell debris. The supernatant was applied onto a self-packaged Ni-affinity column (2 mL Ni-NTA, Genscript) and contaminant proteins were removed with wash buffer (50 mM Tris pH 8.0, 300 mM NaCl, 30 mM imidazole). Then the protein was eluted with elute buffer (50 mM Tris pH 8.0, 300 mM NaCl, 300 mM imidazole). The eluant of protein was concentrated and further purified using a Superdex-200 increase 10/300 GL (GE Healthcare) column equilibrated with a buffer containing 10 mM Tris-HCl pH 8.0, 200 mM NaCl and 5 mM DTT. The purified proteins were analyzed by SDS-PAGE. The fractions containing the target protein were pooled and concentrated. Specifically, SBS Tad1 was purified in the same approach as above, but the buffer pH was 8.8, with 500 mM NaCl and 10% glycerol throughout the whole purification process.

The cells expressing CapV were resuspended with lysis buffer containing 50 mM phosphate buffer pH 7.4, 300 mM NaCl, 10% glycerol (v/v). The CapV proteins bound to Ni-NTA beads were washed with a buffer containing 50 mM phosphate buffer pH 7.4, 300 mM NaCl, 10% glycerol (v/v), 30 mM imidazole and then eluted with the 50 mM phosphate buffer pH 7.4, 300 mM NaCl, 10% glycerol (v/v), 300 mM imidazole. The eluant of CapV was concentrated and further purified using a Superdex-200 increase 10/300 GL (GE Healthcare) column equilibrated with a reaction buffer containing 50 mM phosphate buffer pH 7.4, 300 mM NaCl, 10% glycerol (v/v). The purified protein was analyzed as described above. The fusion protein of NucC with His6-SUMO tag was digested with Ulp1 on the Ni-NTA column at 18°C for 2 h after removing contaminant proteins with wash buffer. Then the NucC protein was eluted with wash buffer. The eluant of NucC was concentrated and further purified as His-tagged proteins as described above.

The full-length HgmTad2 and AcrIIA11 gene was synthesized by GenScript and amplified by PCR and cloned into pGEX6p-1 to produce a GST-tagged fusion protein with a PreScission Protease cleavage site between GST and the target protein. The HgmTad2 mutants were subcloned, overexpressed and purified in the same way as wild-type protein. The proteins were expressed and induced similarly as above. After growth at 16℃ for 12 h, the cells were harvested, re-suspended in lysis buffer (1×PBS, 2 mM DTT and 1 mM PMSF) and lysed by sonication. The cell lysate was centrifuged at 18,000 g for 50 min at 4℃ to remove cell debris. The supernatant was applied onto a self-packaged GST-affinity column (2 mL glutathione Sepharose 4B; GE Healthcare) and contaminant proteins were removed with wash buffer (1×PBS, 2 mM DTT). The fusion protein was then digested with PreScission protease at 16℃ for 2 hours. The protein with an additional five-amino-acid tag (GPLGS) at the N-terminus was eluted with buffer containing 25 mM HEPES pH 7.5, 200 mM NaCl, and 2 mM DTT. The eluant was concentrated and further purified using a Superdex-200 (GE Healthcare) column equilibrated with a buffer containing 10 mM Tris–HCl pH 8.0, 200 mM NaCl, and 5 mM DTT. And then, the HgmTad2 protein was desalted into QA buffer containing 25 mM Tris pH 8.0, 10 mM NaCl and 2 mM DTT by a desalting column (GE Healthcare), and was further purified by ion exchange chromatography with Resource Q column (GE Healthcare). The protein bound to the column was eluted with a gradient concentration of 10-100 mM NaCl, and then protein purity and states were verified with native PAGE and SDS-PAGE, respectively, together with the protein sample flowed through the column. Selenomethionine (Se-Met)-labelled HgmTad2 was expressed in *E. coli* B834 (DE3) cells grown in M9 minimal medium supplemented with 60 mg/L SeMet (Acros) and specific amino acids: Ile, Leu and Val at 50 mg/L; Lys, Phe and Thr at 100 mg/L. The SeMet protein was purified as described above. The four Acb2 homologs were cloned and purified as described previously ^27^.

### Crystallization

All the protein samples in this study were diluted in buffer containing 10 mM Tris-HCl pH 8.0, 200 mM NaCl and 5 mM DTT before crystallization. Each protein was crystallized at 18°C using the following conditions:

#### (1) apo CmTad1/HgmTad2

The concentration of both proteins was 30 mg/mL. The crystals of CmTad1 were grown for 3-4 days using reservoir solution containing 2.0 M ammonium sulfate, 0.1 M sodium HEPES pH 7.5 and 1.4% v/v PEG 400. The crystals of HgmTad2 were grown for 2-3 days using reservoir solution containing 1.0 M lithium chloride, 0.1 M citrate pH 4.0, 20% w/v PEG 6000. Before being harvested, the crystals were cryoprotected in the reservoir solution containing 20% glycerol before flash-freezing in liquid nitrogen.

#### (2) CmTad1 complexed with cA_3_/cAAG, CbTad1 complexed with cA_3_/2’,3’-cGAMP+cA_3_

Prior to crystallization, cA_3_ or cAAG was mixed with protein at a molar ratio of 0.8:1, and 2’,3’-cGAMP was mixed with protein at a molar ratio of 1.2:1. The crystals of CmTad1-cA_3_ and CmTad1-cAAG grew to full size in about 4-5 days, their reservoir solution contains 1.6 M ammonium sulfate, 10% v/v 1,4-Dioxane. The crystallization condition of CbTad1-cA_3_ was 0.1 M MIB (sodium malonate dibasic monohydrate, imidazole, boric acid) pH 6.0, 55% v/v MPD, and the crystallization condition of the CbTad1-cA_3_ was 0.1 M PCTP (sodium propionate, sodium cacodylate trihydrate, Bis-Tris propane) pH 8.0, 60% MPD.

#### (3) CbTad1 complexed with 1′′-3′ gcADPR/1′′-3′ gcADPR+cA_3_

CbTad1 co-expressed with ThsB’ was purified, and then was mixed with cA_3_ at a molar ratio of 1: 0.8. The crystals of purified CbTad1 or its mix with cA_3_ was grown in reservoir solution containing 3.2 M ammonium sulfate and 0.1 M citrate pH 5.0 for 4-5 days. They were stored in antifreeze containing 20% glycerol and quick frozen with liquid nitrogen.

#### (4) HgmTad2 complexed with cGG/1′′-2′ gcADPR/1′′-2′ gcADPR+cGG/3’,3’-cGAMP

These four structures were crystallized in the same condition containing 1.0 M lithium chloride,0.1 M Citrate pH 4.0, 20% w/v PEG 6000. For HgmTad2 complexed with cGG, purified HgmTad2 in the cGG-bound state was used. For HgmTad2 complexed with 1′′-2′ gcADPR or 1′′-2′ gcADPR+cGG, HgmTad2 co-expressed with BdTIR was purified, and no-cGG or cGG-bound form was used, respectively. For HgmTad2 complexed with 3’,3’-cGAMP, HgmTad2 in the no-cGG state was used and mixed with 3’,3’-cGAMP at a molar ratio of 1:1.2.

#### (5) HgmTad2 complexed with 1′′-3′ gcADPR+cGG

HgmTad2 co-expressed with ThsB’ was purified and the cGG-bound form was used in crystallization. The crystallization condition was 0.2 M ammonium sulfate, 0.1 M sodium acetate trihydrate pH 4.6, 25% w/v PEG 4000.

#### (6) apo SPO1 Tad2

After purifying SPO1 Tad2, the protein was diluted to 24 mg/mL, and then grown under the conditions that 0.5 M ammonium sulfate, 1.0 M sodium citrate tribasic dihydrate pH 5.6, 1.0 M lithium sulfate monohydrate conditions for about 1 week.

#### (7) SptTad2-cGG

The SptTad2 protein purified from *E. coli* Bl-21 naturally carries c-di-GMP. The protein was diluted to 24 mg/mL, and then grown under the condition containing 0.3 M magnesium nitrate hexahydrate,0.1 M Tris pH 8.0, 23% w/v PEG2000 for 4-5 days, and then transferred into antifreeze and then flash-freezing in liquid nitrogen.

### Data collection, structure determination and refinement

All the data were collected at SSRF beamlines BL02U1 and BL19U1, integrated and scaled using the HKL2000 package ^41^. The initial model of CbTad1 was used from PDB: 7UAV. The initial models of CmTad1, SPO Tad2 and SptTad2 were obtained using AlphaFold2 ^42^. The structure of apo HgmTad2 was solved by SAD phasing, using Autosol in PHENIX ^43^. The structures of protein complexed with cyclic oligonucleotides were solved through molecular replacement and refined manually using COOT ^44^. All the structures were further refined with PHENIX ^43^ using non-crystallographic symmetry and stereochemistry information as restraints. The final structure was obtained through several rounds of refinement. Final Ramachandran statistics: 96.75% favoured, 3.25% allowed and 0% outliers for apo-CmTad1-Zn structure; 96.62% favoured, 3.38% allowed and 0% outliers for CmTad1-Zn-cA_3_; 96.75% favoured, 3.25% allowed and 0% outliers for CmTad1-Zn-cAAG structure; 97.59% favoured, 2.41% allowed and 0% outliers for CbTad1-1’’,3’-gcADPR structure; 96.37% favoured, 3.63% allowed and 0% outliers for CbTad1-1’’,3’-gcADPR-cA_3_ structure; 96.99% favoured, 3.01% allowed and 0% outliers for CbTad1-2’,3’-cGAMP structure; 97.82% favoured, 2.18% allowed and 0% outliers for CbTad1-2’,3’-cGAMP-cA_3_ structure; 97.55% favoured, 2.45% allowed and 0% outliers for apo-HgmTad2 structure; 96.32% favoured, 3.68% allowed and 0% outliers for HgmTad2-1’’,2’-gcADPR structure; 95.34% favoured, 4.66% allowed and 0% outliers for HgmTad2-1’’,2’-gcADPR-cGG structure; 97.06% favoured, 2.94% allowed and 0% outliers for HgmTad2-1′′-3′-gcADPR-cGG structure; 97.06% favoured, 2.94% allowed and 0% outliers for HgmTad2-cGG structure; 98.77% favoured, 1.23% allowed and 0% outliers for HgmTad2-3’,3’-cGAMP structure; 97.98% favoured, 2.02% allowed and 0% outliers for SptTad2-cGG structure; 98.63% favoured, 1.37 allowed and 0% outliers for apo-SPO1 Tad2 structure. Structural illustrations were generated using PyMOL (https://pymol.org/). Data collection and structure refinement statistics are summarized in Extended Data Table 1.

### Isothermal titration calorimetry binding assay

The dissociation constants of binding reactions of CmTad1/CbTad1 with cA_3_/cAAG/3’,2’-cGAMP/2’,3’-cGAMP/3’,3’-cGAMP/cAA/cGG/cUG/cUA/cUU, SBS Tad1 with cA_3_/cAAG/3’,2’-cGAMP/2’,3’-cGAMP/3’,3’-cGAMP/cUA/1′′–2′ gcADPR/1′′–3′ gcADPR, and ColiTad1 with cA_3_/cAAG/3’,2’-cGAMP/2’,3’-cGAMP/3’,3’-cGAMP were determined by isothermal titration calorimetry (ITC) using a MicroCal ITC200 calorimeter. All the protein and cyclic-oligonucleotides were desalted into the working buffer containing 20 mM HEPES pH 7.5 and 200 mM NaCl. The titration, for example, was carried out with 19 successive injections of 2 µL cA_3_/cAAG at 25 µM concentration, spaced 120 s apart, into the sample cell containing CbTad1 with a concentration of 5 µM by 700 rpm at 25°C. Correspondingly, 3’,2’-cGAMP/2’,3’-cGAMP at 150 µM concentration was titrated into 50 µM CbTad1. cA_3_/cAAG at 150 μM concentration was titrated into 30 μM CmTad1, and 3’,2’-cGAMP/2’,3’-cGAMP/3’,3’-cGAMP/cAA/cGG/cUG/cUA/cUU at 300 μM concentration was titrated into 100 μM CmTad1. For SBS Tad1, 3’,2’-cGAMP/3’,3’-cGAMP/2’,3’-cGAMP/cUA/1′′–2′ gcADPR/1′′–3′gcADPR at 300 μM concentration was titrated into 100 μM SBS Tad1, and 3’,2’-cGAMP/3’,3’-cGAMP/2’,3’-cGAMP at 300 μM concentration was titrated into 100 μM ColiTad1. For both SBS Tad1 and ColiTad1, cA_3_/cAAG at 150 μM concentration was titrated into 30 μM SBS Tad1 or ColiTad1. All of the above titration experiments were performed in the same experimental procedure. The Origin software was used for baseline correction, integration, and curve fitting to a single site binding model.

### ThsA NADase activity assay

NADase assay was performed by using ThsA enzyme from *Bacillus cereus* MSX-D12, which was expressed and purified as described previously, as a reporter for the presence of cyclic ADPR isomers. NADase reaction was performed in a black, 96-well plate (Corning 96-well half area black non-treated plate with a flat bottom) at 37 °C in a 95 µL reaction volume, and the final concentration of ThsA and 1′′–3′ gcADPR was 50 and 5 nM, respectively. Next, 5 µL of 2 mM ε-NAD solution was added to each well immediately before measurement and mixed by pipetting rapidly. ε-NAD was used as a fluorogenic substrate to report ThsA enzyme NADase activity by monitoring increase in fluorescence (excitation 300 nm, emission 410 nm) using EnSpire Multimode Plate Reader (PerkinElmer) at 37 °C. To examine the inhibitory effect of HgmTad2 or its mutants on ThsA, HgmTad2 or its mutants (40 nM of each) was incubated with 5 nM 1′′–3′ gcADPR in incubation buffer (50 mM Tris pH 7.5 and 50 mM NaCl) at room temperature for 5 min in advance. Then ThsA was added at a final concentration of 50 nM. After an incubation for 5 minutes, ε-NAD was added to start the reaction.

### Surface Plasmon Resonance assay

The SPR analysis was performed using a Biacore 8K (GE Healthcare) at room temperature (25 ℃). Equal concentrations of HgmTad2/SPO1 Tad2/SptTad2/SaTad2 were immobilized on channels of the carboxymethyldextran-modified (CM5) sensor chip to about 280 Response Unit (RU). To collect data for kinetic analysis, a series of concentrations (12.5 nM, 25 nM, 50nM, 100 nM, 200 nM) of 3’,3’-cGAMP/3’,2’-cGAMP/2’,3’-cGAMP/cGG/cUG/cA_3_ diluted in binding buffer (20 mM HEPES pH 7.5, 200 mM NaCl and 0.05% (v/v) Tween-20) was injected over the chip at a flow rate of 30 μL/min. The protein-ligand complex was allowed to associate for 60 s and dissociate for 600 s. Data were fit with a model describing a bivalent analyte. Kinetic rate constants were extracted from this curve fit using Biacore evaluation software (GE healthcare).

### High-performance liquid chromatography (HPLC)

For analysis of ligand sequestering, 40 μM Tad1 or Tad2 protein was pre-incubated with 4 μM cA_3_, 2’,3’-cGAMP or 3’,3’-cGAMP for 30 min at 18°C. And then, for Tad1 series, proteinase K was subsequently added to the reaction system at a final concentration of 0.5 μM and the reaction was performed at 58°C for 1 h. For Tad2, the sample was first heated at 100°C for 10 min, and then proteinase K was subsequently added to the reaction system at a final concentration of 25 μM and the reaction was performed at 58°C for 3 h. For analysis of intrinsically bound nucleotide in HgmTad2 during expression, 40 μM HgmTad2 in different states was treated as reported for Tad2 in the above. 4 μM 3’,3’-cGAMP and cGG were used as standards.

Reaction samples were transferred to Amicon Ultra-15 Centrifugal Filter Unit 3 kDa and centrifuged at 4°C, 4,000 g. The products obtained by filtration were further filtered with a 0.22 μm filter and subsequently used for HPLC experiments. The HPLC analysis was performed on an Agilent 1200 system with a ZORBAX Bonus-RP column (4.6 × 150 mm). A mixture of 2% acetonitrile and 0.1% trifluoroacetic acid solution in water (98%) were used as mobile phase with 0.8 mL/min. For cA_3_, 5% acetonitrile and 0.1% trifluoroacetic acid solution in water (95%) were used as mobile phase. The compounds were detected at 254 nm.

### Fluorogenic biochemical assay for CapV activity

The enzymatic reaction velocity was measured as previously described ^18^. Briefly, the esterase activity of the 6×His-tagged CapV was probed with the fluorogenic substrate resorufin butyrate. The CapV protein was diluted in 50 mM sodium phosphate pH 7.4, 300 mM NaCl, 10% (v/v) glycerol to a final concentration of 2 μM. To determine the enzymatic activity of CapV activated by 3’,3’-cGAMP, 0.8 μM of 3’,3’-cGAMP was added to DMSO solubilized resorufin butyrate (stock of 20 mM mixed with 50 mM sodium phosphate pH 7.4, 300 mM NaCl, 10% v/v glycerol reaching a final concentration of 100 μM). Subsequently, the purified 6×His-tagged CapV was added to the reaction solution containing 3’,3’-cGAMP to a final assay volume of 50 μL, and fluorescence was measured in a 96-well plate (Corning 96-well half area black non-treated plate with a flat bottom). Plates were read once every 30 s for 10 min at 37°C using a EnSpire Multimode Plate Reader (PerkinElmer) with excitation and emission wavelengths of 550 and 591 nm, respectively.

To determine the function of inhibitory proteins, 8 μM protein was pre-incubated with 0.8 μM 3’,3’-cGAMP for 10 min at 18°C, and the subsequent detection method was as described above. To examine whether the released molecule from HgmTad2 or SBS Tad1 is still able to activate CapV, 0.8 μM 3’,3’-cGAMP was incubated with 8 μM HgmTad2 or SBS Tad1 for 10 min at 18°C. Proteinase K was subsequently added to the reaction system at a final concentration of 25 μM and the reaction was performed at 58°C for 3 h. Reaction products were transferred to Amicon Ultra-4 Centrifugal Filter Unit 3 kDa and centrifuged at 4°C, 4,000 g. Filtered products were used for CapV activity assay as described above.

### *Sf*TIR-STING NAD_+_ cleavage activity analysis

The enzymatic reaction velocity was measured as previously described ^45^. The enzymatic activity of *Sf*TIR-STING was activated by cGG. 500 μM ε-NAD and 50 nM cGG were mixed in a 96-well plate format with reaction buffer (50 mM Tris pH 7.5, 50 mM NaCl). Subsequently, purified 6×His-tagged *Sf*TIR-STING was added to the reaction to a final assay volume of 50 μL and plates were read once every 15 s for 10 min at 37°C using a EnSpire Multimode Plate Reader (PerkinElmer) with excitation and emission wavelengths of 410 and 300 nm, respectively. Reaction rate was calculated from the linear part of the initial reaction.

To determine the function of inhibitory proteins, 1 μM HgmTad2, SPO1 Tad2, SptTad2 and SaTad2 was pre-incubated with 50 nM cGG for 20 min at 18°C, and the subsequent detection method was as described above. To determine the function of HgmTad2 mutants, 200 nM HgmTad2 and its mutants was pre-incubated with 50 nM cGG for 20 min at 18°C. To examine whether the released molecule from HgmTad2 is still able to activate *Sf*TIR-STING, 50 nM cGG was incubated with 200 nM HgmTad2 for 10 min at 18°C. Proteinase K was subsequently added to the reaction system at a final concentration of 1 μM and the reaction was performed at 58°C for 1 h. Reaction products were transferred to Amicon Ultra-4 Centrifugal Filter Unit 3 kDa and centrifuged at 4°C, 4,000 g. Filtered products were used for *Sf*STING activity assay as described above.

### *In vitro* NucC activity assay

The nuclease activity assay was measured as previously described ^27^. Plasmid pUC19 was used as substrate. NucC (10 nM) and cA_3_ molecules (5 nM) were mixed with 0.4 μg DNA in a buffer containing 25 mM Tris-HCl pH 8.0, 10 mM NaCl, 10 mM MgCl_2_, and 2 mM DTT (20 μL reaction volume), incubated at 37°C for 20 min, then separated on a 1% agarose gel. Gels were stained with Goldview and imaged by UV illumination.

To determine the function of CbTad1, 200 nM CbTad1 or its mutants were pre-incubated with the other components at 18°C for 30 min, and the subsequent reaction and detection method was as described above. To examine whether the released molecule from CbTad1 is able to activate NucC, 5 nM cA_3_ was incubated with 200 nM CbTad1 at 18°C for 20 min. Proteinase K was subsequently added to the reaction system at a final concentration of 1 μM and the reaction was performed at 58°C for 1 h, then the proteinase K-treated samples were heated with 100°C for 10 min to extinguish proteinase K and the subsequent detection method was as described above.

### *In vitro* SpyCas9 DNA cleavage assay

SpyCas9 sgRNA was generated by *in vitro* T7 transcription kit (Vazyme). 100 nM SpyCas9 and 150 nM sgRNA was incubated with 10 μM purified Tad2 or AcrIIA11 in cleavage buffer (20 mM HEPES-KOH pH 7.5, 75 mM KCl, 10% glycerol, 1 mM DTT, and 10 mM MgCl_2_) for 30 min at 37°C. Plasmid pUC57 containing the target protospacer 25 sequence inserted using *KpnI/XbaI* was linearized by ScaI digestion. Linearized plasmid was added to the Cas9/sgRNA complex at 10 nM final concentration. The reactions were incubated at 37°C for 10 min and extinguish by 1 μM proteinase K for 15 min at 58°C, then separated on a 1% agarose gel. Gels were stained with Goldview and imaged by UV illumination.

#### SpyCas9_sgRNA_DNA template

ATGTAATACGACTCACTATAGGAAATTAGGTGCGCTTGGCGTTTTAGAGCTAGAAATAG CAAGTTAAAATAAGGCTAGTCCGTTATCAACTTGAAAAAGTGGCACCGAGTCGGTGCTT

#### Cleavage assay DNA sequence

TCGGTGCGGGCCTCTTCGCTATTACGCCAGCTGGCGAAAGGGGGATGTGCTGCAAGGCGATTAAGTTGGGTAACGCCAGGGTTTTCCCAGTCACGACGTTGTAAAACGACGGCCAGTGCCAAGCTTGCATGCCTGCAGGTCGACTCTAGAGGATCCCAATCCCAGCCAAGCGCACCTAATTTCCGAATTCGTAATCATGGTCATAGCTGTTTCCTGTGTGAAATTGTTATCCGCTCACAATTCCACACAACATACGAGCCGGAAGCATAAA

### Native-PAGE assay

For ligand binding native-PAGE assay, proteins were pre-incubated with cyclic nucleotides for 20 min at 18°C, where protein was 20 μM and the concentrations of cyclic nucleotides was 5, 10 or 20 μM, respectively. Products of the reaction were analyzed using 5% native polyacrylamide gels and visualized by Coomassie blue staining.

### Multi-angle light scattering (MALS)

MALS experiments were performed in 10 mM Tris pH 8.0, 200 mM NaCl and 2 mM TCEP using a Superdex-200 10/300 GL size-exclusion column from GE Healthcare. All protein concentrations were diluted to 1.7 mg/mL. The chromatography system was connected to a Wyatt DAWN HELEOS Laser photometer and a Wyatt Optilab T-rEX differential refractometer. Wyatt ASTRA 7.3.2 software was used for data analysis.

### Episomal gene expression

The shuttle vector pHERD30T that replicates in *P. aeruginosa* and *E. coli* ^46^ was used for episomal expression of Acb2 and Tad proteins in *P. aeruginosa* strains. pHERD30T has an arabinose-inducible promoter and a selectable gentamicin marker. Vector was digested with NcoI and HindIII restriction enzymes. Inserts were amplified by PCR using bacterial overnight culture or synthesized by Twist Bioscience and joined with the digested vector using Hi-Fi DNA Gibson Assembly (NEB) following the manufacturer’s protocol. The resulting plasmids were transformed into *E. coli* DH5ɑ. All plasmid constructs were verified by whole plasmid sequencing. *P. aeruginosa* cells were electroporated with the pHERD30T constructs and selected on gentamicin.

### Chromosomal Thoeris integration

For chromosomal insertion of the MRSN390231 Thoeris SIR2 (Pa231) operon at the Tn7 locus in *P. aeruginosa* PAO1(PAO1 Tn7:Thoeris SIR2), the integrating vector pUC18-mini-Tn7T-LAC ^47^ carrying Thoeris operon and the transposase expressing helper plasmid pTNS3 ^48^ were used. pUC18-mini-Tn7T-LAC empty vector was used for the creation of the control strain (PAO1 Tn7:empty). The vector was linearized using around-the-world PCR (in positions of KpnI and BamHI sites), treated with DpnI, and then purified. The insert was amplified using MRSN390231 overnight culture as a DNA template and joined with linearized pUC18-mini-Tn7T-LAC vector using Hi-Fi DNA Gibson Assembly (NEB) following the manufacturer’s protocol. The resulting plasmids were used to transform *E. coli* DH5ɑ. All plasmid constructs were verified by whole plasmid sequencing. *P. aeruginosa* PAO1 cells were electroporated with pUC18-mini-Tn7T-LAC and pTNS3 and selected on gentamicin-containing plates. Potential integrants were screened by colony PCR with primers PTn7R and PglmS-down ^48^. Electrocompetent cell preparations, transformations, integrations, selections, plasmid curing, and FLP-recombinase-mediated marker excision with pFLP were performed as described previously ^49^.

### Phage growth

Phages F10 and JBD67Δ*acb2* were grown on *P. aeruginosa* PAO1, which lacks CBASS and Thoeris systems. Phage PaMx41Δ*acb2* was grown on *P. aeruginosa* BWHPSA011 (Pa011) ΔCBASS strain. For phage propagation 100 µl of *P. aeruginosa* overnight cultures were infected with 10 µl of low titer phage lysate (>10^4–7^ pfu/ml) and then mixed with 3 ml of 0.35% top agar 10 mM MgSO_4_ for plating on the LB solid agar (20 ml LB agar with 10 mM MgSO_4_). After incubating 37 °C overnight, 2.5 ml SM phage buffer was added on the solid agar lawn and then incubated for 10 minutes at room temperature. The whole cell lysate was collected, a 10% volume of chloroform was added, and the tubes were left for 20 minutes at room temperature with gentle shaking, followed by centrifugation at maximum speed for 3 min 4°C to remove cell debris. The supernatant phage lysate was stored at 4°C for downstream assays.

### Plaque assays

Plaque assays were conducted at 37 °C with solid LB agar plates supplemented with 10 mM MgSO_4,_ 50 µg ml^-1^ gentamicin, 0.2% L-arabinose, and 0.3 mM IPTG for PAO1 strains with CBASS or Thoeris operon chromosomal integration, and the same conditions except without IPTG for the native CBASS and Thoeris strains. 100 μL of overnight bacterial culture was mixed with top agar (0.35% agar in LB) and plated. Phage lysates were diluted 10-fold then 2.5 μL spots were applied to the top agar after it had been poured and solidified. The plates were incubated overnight at 37 °C.

### Bioinformatic analysis of CD-NTases

CD-NTases were identified within the bacterial hosts relevant for each Tad protein by using a protein BLAST (blastp) search. A previously curated list of CD-NTase sequences ^21^was queried against *Clostridium* (taxid:1485), *Clostridoioides* (taxid:1870884), *Bacteroides* (taxid:816), *Sphingobacterium* (taxid:28453), and *Bacillus cereus* group (taxid:86661). There is only one CD-NTase record for *Salegentibacter* (taxid:143222) and zero for *Colidextribacter* (taxid:1980681), so a list of CD-NTase across bacterial taxonomies was used from Whiteley et al. 2019. These lists of CD-NTases were queried against the NCBI non-redundant protein database of each respective bacterial genus as indicated above. A genus-level analysis was chosen due to the diversity of CD-NTase sequences associated with the different clades, which are known or predicted to produced specific cyclic oligonucleotides. Hits from the blastp search with >24.5% amino acid identity, >50% coverage, and an E value of <0.0005 were identified as CD-NTases. Two or more CD-NTases per CD-NTase clade per bacterial genus were queried using Defense Finder ^50,51^, which revealed that all CD-NTases identified are a part of a CBASS system. 183 CD-NTase hits were identified in *Clostridium* and nine hits in *Clostridoides*, so their results were combined as 202 total hits in Extended Data Figure 3. A total of 107 hits were identified in *Bacteroides*, 71 in *Sphingobacterium*, and 270 in *Bacillus cereus group*. Six hits were found in *Salegentibacter* and zero hits for *Colidextribacter*.

### Phylogenetic tree analysis

Tad1 and Tad2 homologs were identified using SBSTad1 (NCBI: P0DW57) and SPO1Tad2 (NCBI: YP_002300464.1), respectively, as query proteins to seed a position-specific iterative blast (PSI-BLAST) search of the NCBI non-redundant protein database. Three rounds of PSI-BLAST searches were performed with a max target sequence of 5,000 and E value cut-off of 0.005 for inclusion in the next search round, BLOSUM62 scoring matrix, gap costs settings existence 11 and extension 1, and using conditional compositional score matrix adjustment. Hits from the third search round of PSI-BLAST with >70% coverage, E value of < 0.0005 and length less than 190 amino acids (for Tad1) and length 70-120 amino acids (for Tad2) were clustered using MMSeq2 ^52^ to remove protein redundancies (minimum sequence identity=0.9 for Tad1 and 0.8 for Tad2, minimum alignment coverage=0.9), which resulted in 410 and 667 representative Tad1 and correspondingly, Tad2 homolog sequences. MAFFT (FFT-NS-I iterative refinement method) ^53^ was used to create protein alignment. Manual analysis of the MAFFT protein alignment was performed to ensure the presence of at least one of the cyclic oligonucleotide binding site regions and to remove non-relevant sequences. The final aligned 385 and 568 sequences (Tad1 and Tad2 correspondingly) were used to construct a phylogenetic tree using FastTree ^54^ and then visualized and annotated in iTOL ^55^.

