## Extended Data Figures and Table for "Single phage proteins sequester TIR- and cGAS-generated signaling molecules"

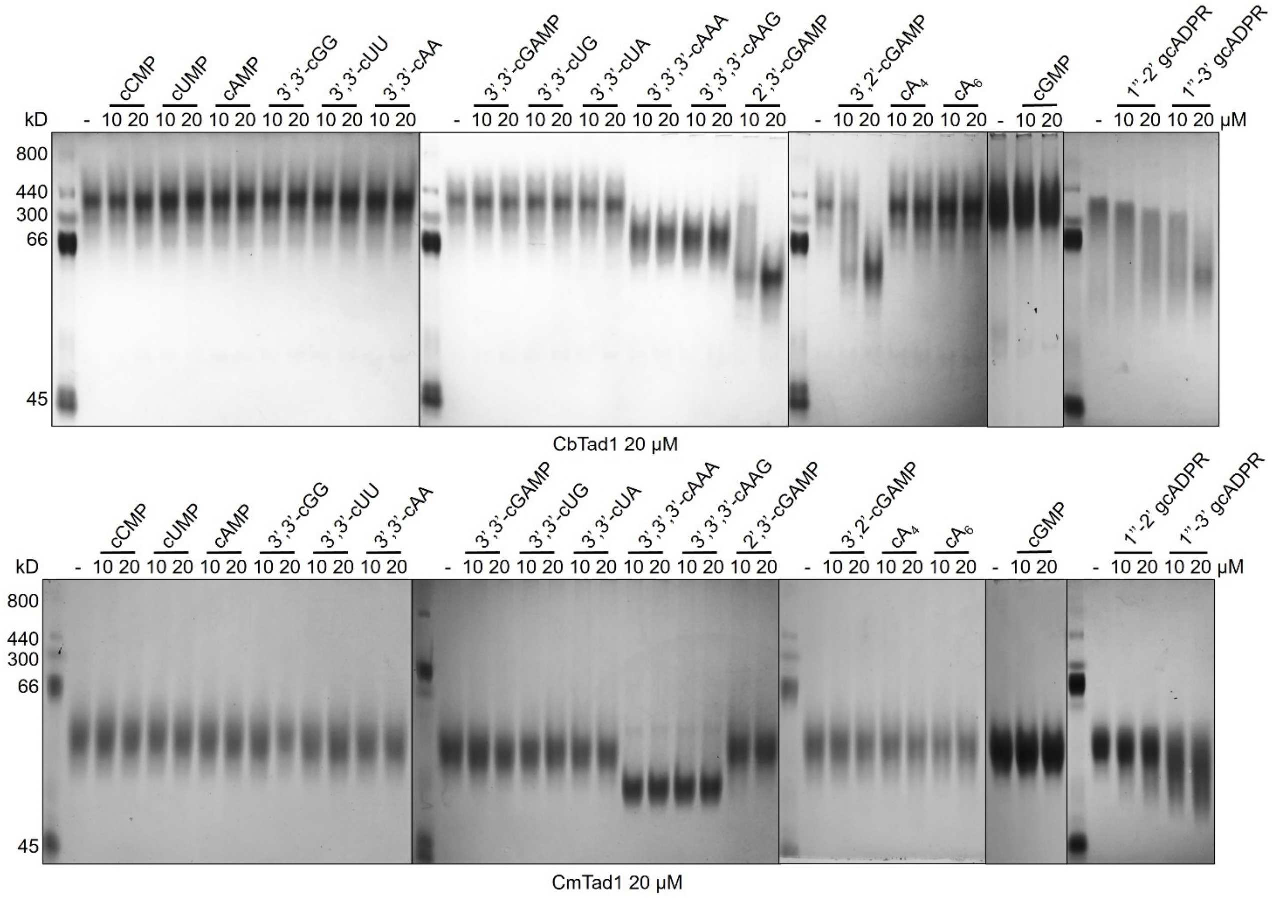


**Extended Data Figure 1. Native gel assay of CbTad1 and CmTad1.**

Native PAGE showed the binding of CbTad1 and CmTad1 to cyclic oligonucleotides and gcADPR molecules.

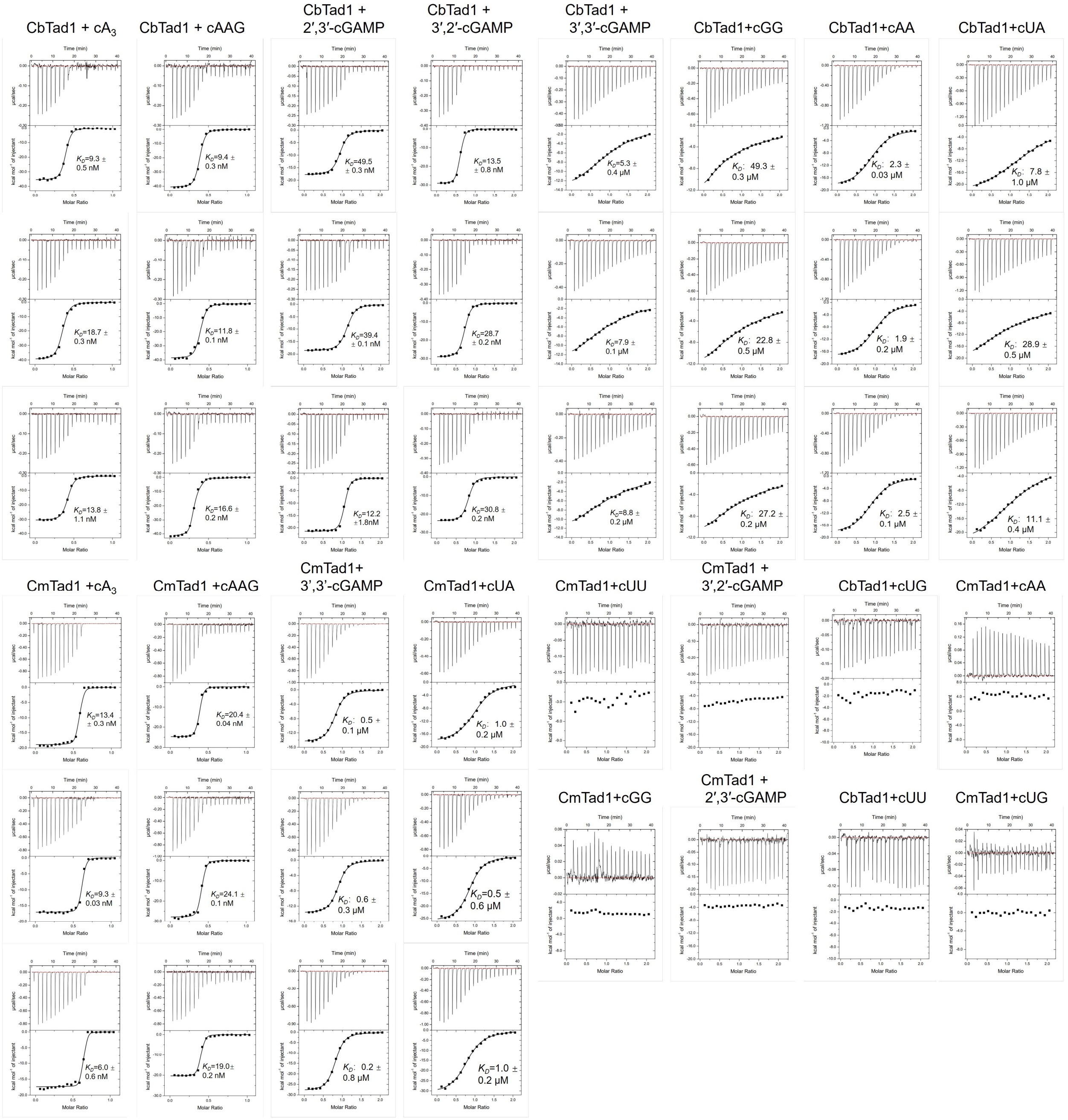


**Extended Data Figure 2. The binding spectrum of CbTad1 and CmTad1 studied by ITC assay.**
ITC assays to test binding of cyclic oligonucleotides to CbTad1 and CmTad1.

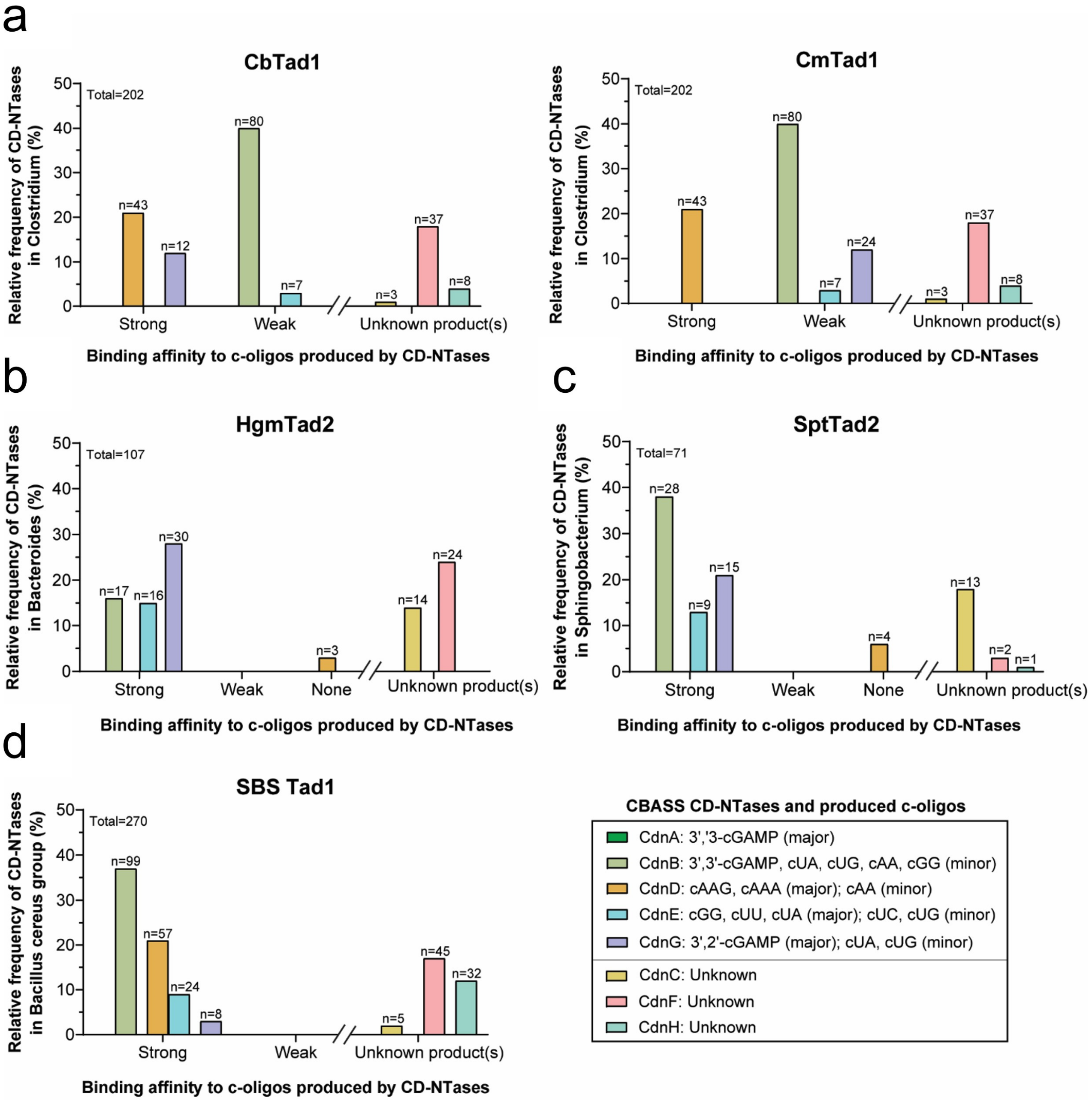


**Extended Data Figure 3. Bacterial hosts that Tad-encoding phage likely infect contain multiple CBASS CD-NTases that produce cyclic oligonucleotides.**

Bacteria from the genus **(a)** *Clostridium*, **(b)** *Bacteroides*, **(c)** *Sphingobacterium*, and **(d)** *Bacillus cereus* group contain CBASS CD-NTases that produce cyclic oligonucleotides (c-oligos) with strong, weak, or no binding affinity to the Tad proteins tested in this study. The relative frequency of CD-NTases is quantified as the number of CD-NTase from a specific clade divided by the total number of CD-NTases identified using the NCBI blastp (see Methods for details). The legend indicates the c-oligos that are known or predicted to be produced by the indicated CBASS CD-NTases (Whiteley et al., 2019; Ye et al., 2020; Morehouse et al., 2020; Fatma et al., 2021). CD-NTases with currently unknown nucleotide products are indicated in the legend and the graphs.

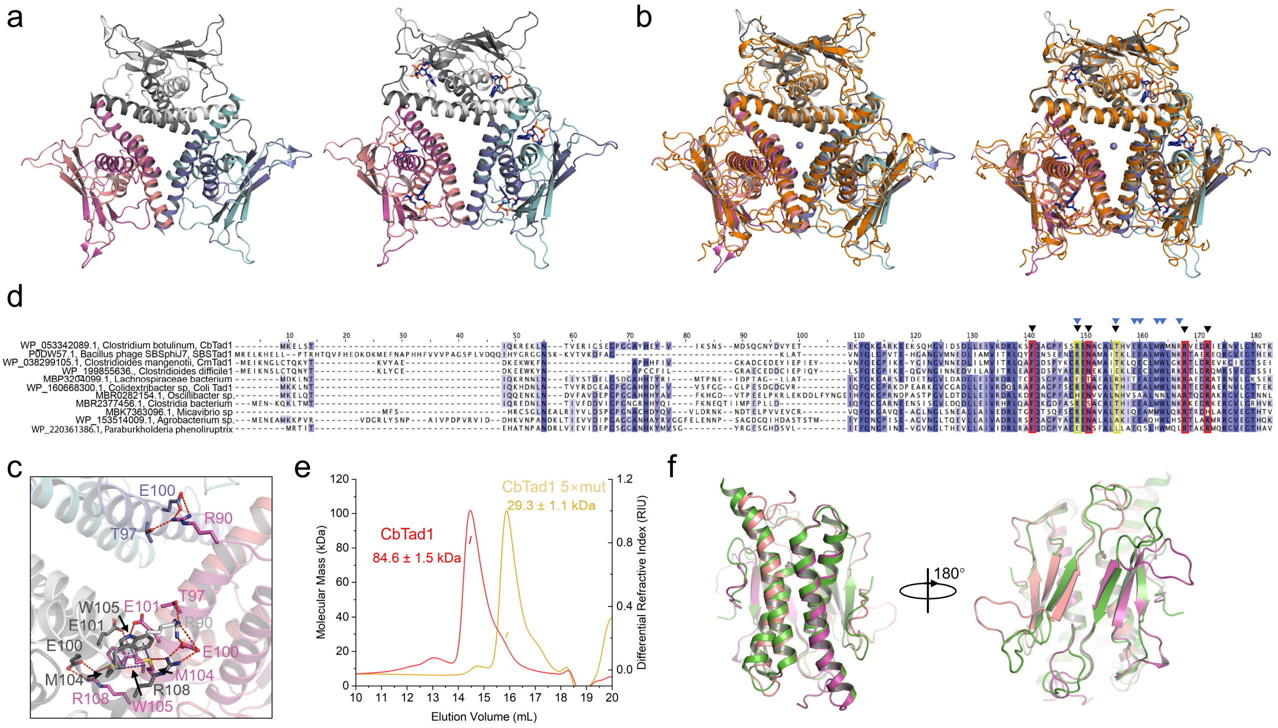


**Extended Data Figure 4. Structural comparison between CbTad1 and CmTad1.**

**a,** The hexameric form of CbTad1 are shown in cartoon model. Left: CbTad1 (PDB code: 7UAV). Right: CbTad1-1''-2' gcADPR (PDB code: 7UAW).

**b,** Structural comparison between CbTad1 and CmTad1 hexamers. Left: structural comparison between apo CbTad1 (colored as in a) and apo CmTad1 (colored orange). Right: structural comparison between CbTad1-1''-2' gcADPR (colored as in a) and apo CmTad1 (colored orange).

**c,** Detailed binding in the hexamer interface of CbTad1. Residues involved in hexamer formation are shown as sticks. Red dashed lines represent polar interactions.

**d,** Sequence alignment between Tad1 homologs. The cyclic trinucleotide (CTN) and cyclic dinucleotides (CDN)/gcADPR binding sites are marked in yellow and red, respectively. Representative sequences were intentionally selected to show CDN/CTN binding site mutations. Residues involved in hexamer formation are marked with blue triangles.

**e,** SLS studies of purified CbTad1 and its E100A/E101A/M104A/W105A/R108A mutant. Calculated molecular weight is shown above the peaks.

**f,** Structural comparison between the dimer form of CbTad1 and CmTad1. Apo CbTad1 (PDB code: 7UAV) is colored green. Apo CmTad1 is colored in pink and magenta for the two protomers.

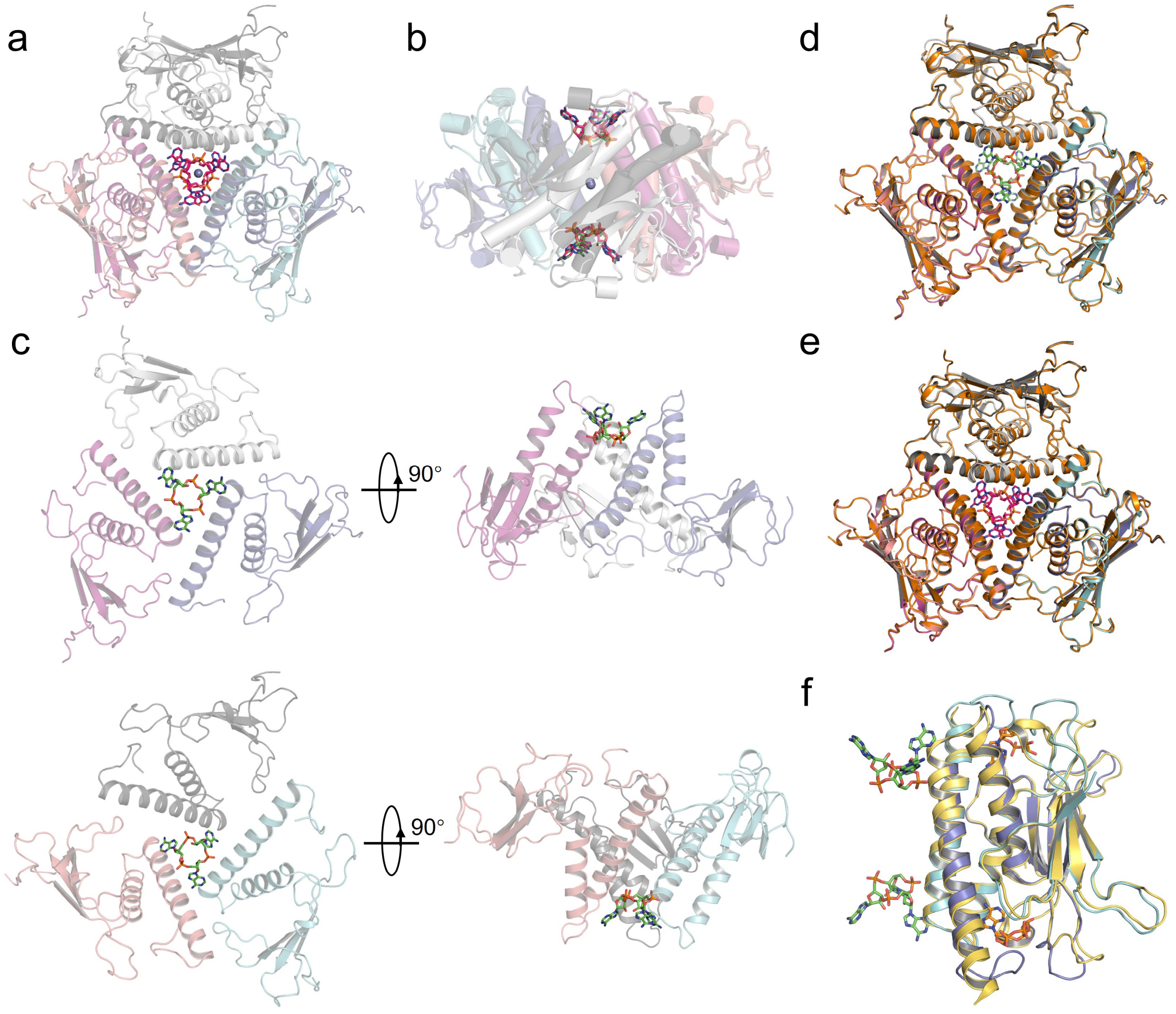


**Extended Data Figure 5. Structures of CmTad1 complexed with $cA_3$ or cAAG.**

**a,** Overall structure of CmTad1 complexed with cAAG.

**b,** Structural comparison between CmTad1-$cA_3$ and CmTad1-cAAG. $cA_3$ and cAAG is colored in green and hot pink sticks, respectively.

**c,** Binding of $cA_3$ by CmTad1. Each $cA_3$ is bound by three protomers of Tad1.

**d, e,** Structural comparison between CmTad1-$cA_3$ (**d**) and apoCmTad1, and between CmTad1-cAAG (**e**) and apoCmTad1. CmTad1-$cA_3$ and CmTad1-cAAG is colored as in Figure 2E and S3A, respectively. Apo CmTad1 is colored in orange.

**f,** Structural alignment between the dimer forms of CbTad1-$cA_3$-1''-3' gcADPR (colored cyan and slate for the two protomers) and apo CbTad1 (colored yellow and orange). $cA_3$ and 1''-3' gcADPR is colored in green and orange sticks, respectively.

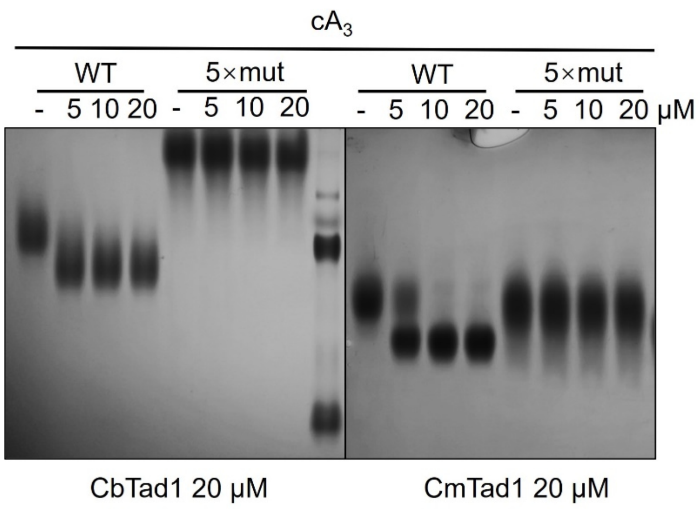


**Extended Data Figure 6. Disruption of Tad1 hexamer abolishes $cA_3$ binding.**
Native PAGE showed the binding of CbTad1 and CmTad1 and their mutants to $cA_3$.

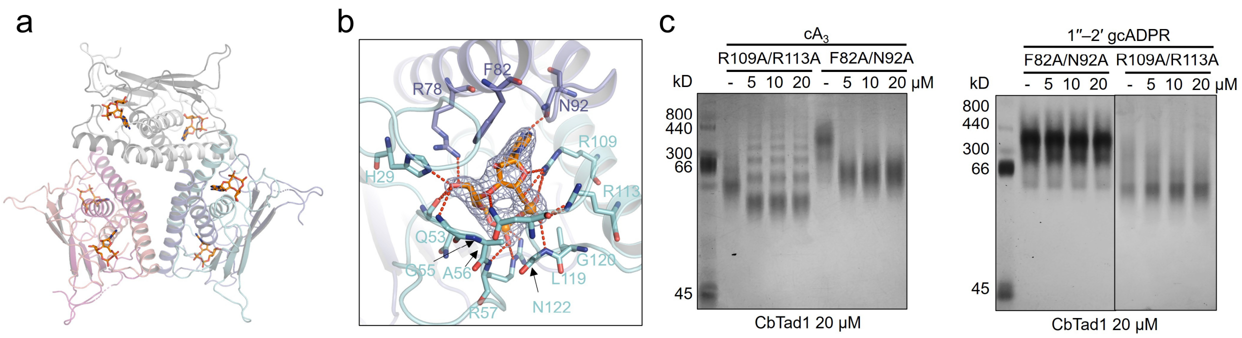


**Extended Data Figure 7. Binding of 1”-3’-gcADPR by CbTad1.**
**a,** Overall structure of CbTad1 hexamer bound to 1”-3’ gcADPR, which is shown as orange sticks.
**b,** Detailed binding between CbTad1 and 1”-3’ gcADPR. Residues involved in 1”-3’ gcADPR binding are shown as sticks. Red dashed lines represent polar interactions. 2Fo-Fc electron density of 1”-3’ gcADPR within one binding pocket is shown and contoured at 1 σ.
**c,** Native PAGE showed the binding of CbTad1 mutants to $cA_3$ and 1”-2’ gcADPR.

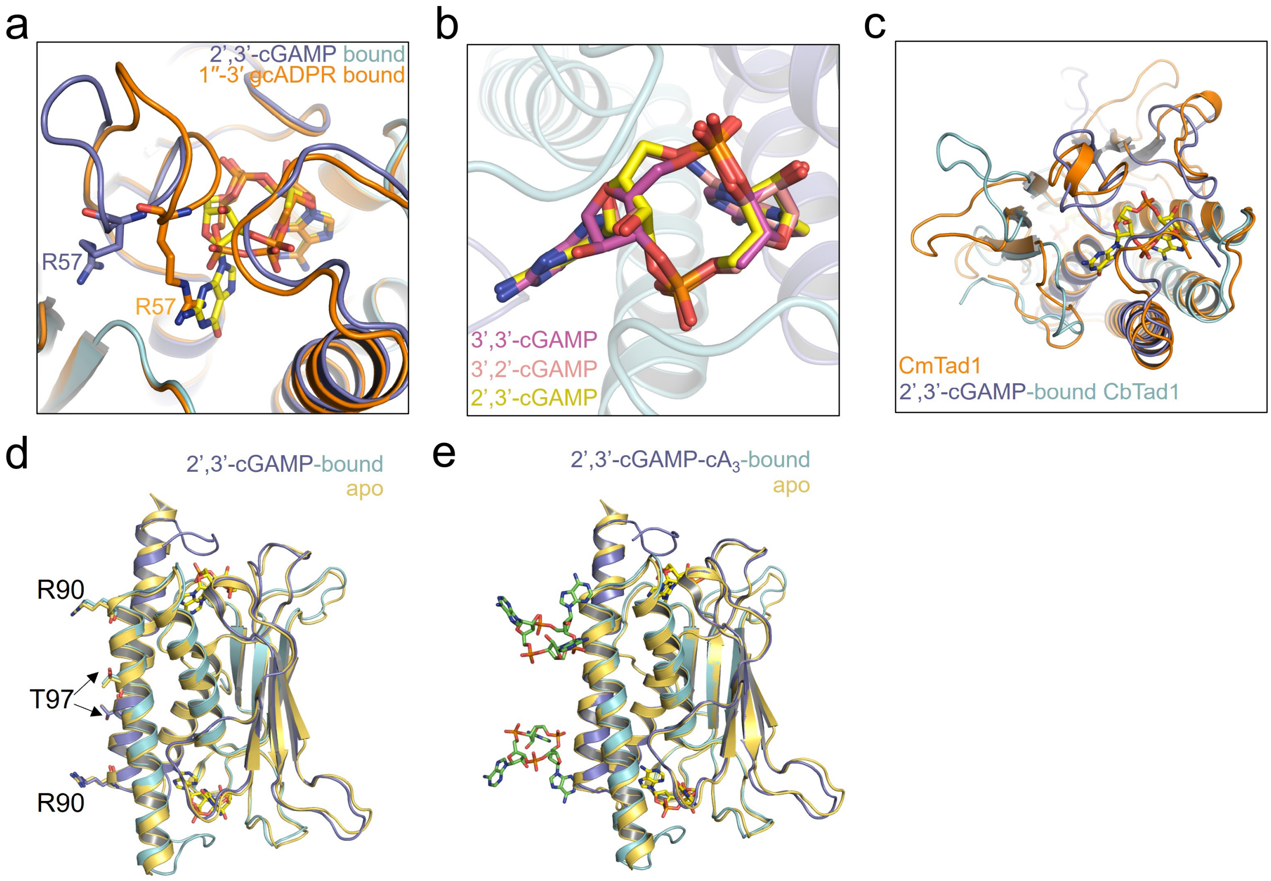


**Extended Data Figure 8. Structural alignments among CbTad1 in apo form and complexed with ligands.**

**a,** Structural alignment between 1''-3' gcADPR-bound (colored orange) and 2',3'-cGAMP-bound CbTad1 (colored slate and cyan for two protomers within a dimer). 2',3'-cGAMP and 1''-3' gcADPR is shown as yellow and orange sticks, respectively. R57 in both structures are shown as sticks.

**b,** Docking of 3',2'-/3',3'-cGAMP into the binding pocket of 2',3'-cGAMP in CbTad1. 3',3'-cGAMP has a backbone structure different from those of 2',3'-/3',2'-cGAMP.

**c,** Structural alignment between 2',3'-cGAMP-bound CbTad1 (colored as in a) and apo CmTad1 (colored orange). 2',3'-cGAMP is shown as yellow sticks.

**d,** Structural alignment between 2',3'-cGAMP-bound CbTad1 and apo CbTad1 (colored yellow). 2',3'-cGAMP-bound CbTad1 is shown as in Figure 3A. Residues involved in binding of cyclic trinucleotides are shown as sticks in both structures.

**e,** Structural alignment between $cA_3$-2',3'-cGAMP-bound CbTad1 and apo CbTad1 (colored yellow). $cA_3$-2',3'-cGAMP-bound CbTad1 is shown as in Figure 3E.

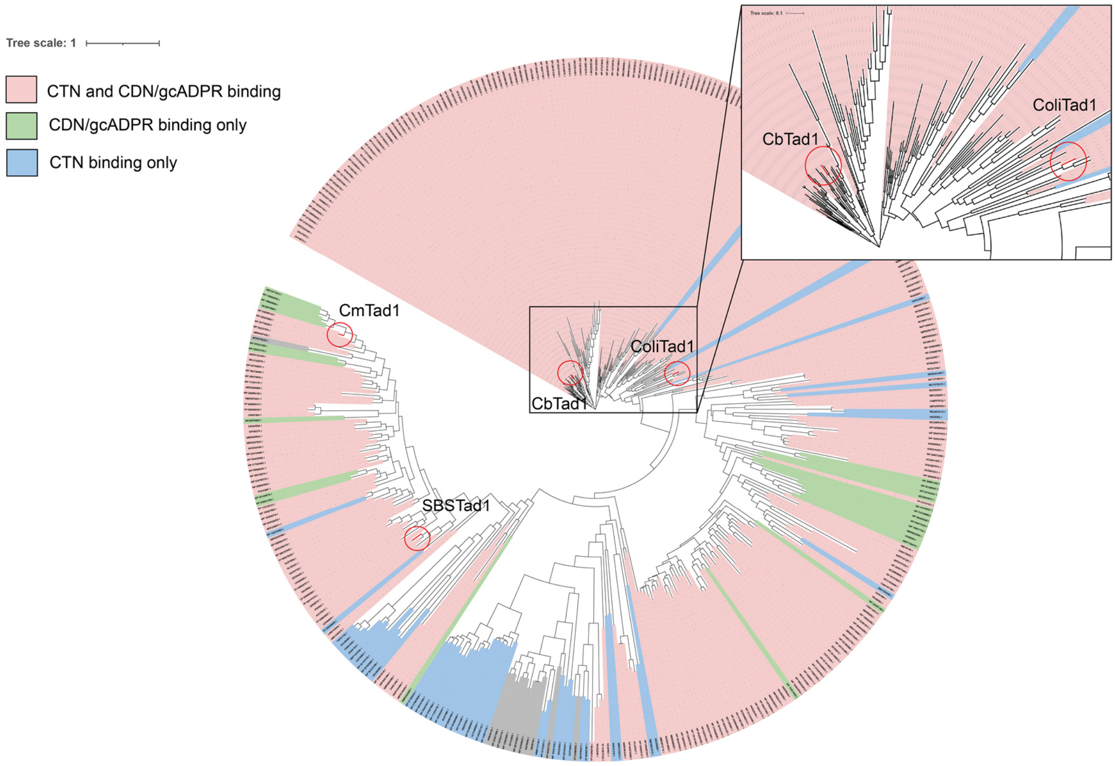


**Extended Data Figure 9. Phylogenetic analysis of Tad1 homologs.**

Phylogenetic analysis of Tad1 homologs found in NCBI non-redundant protein database. The proteins with CTN and CDN/gcADPR binding are predicted to be functional are indicated in red; with substitutions in CTN binding sites (mutant R/T residues) in green; with substitutions in CDN/gcADPR binding sites (mutant F/N or R/R residues) in blue. Tad1 proteins used for biochemical studies are indicated on the tree by red circles.

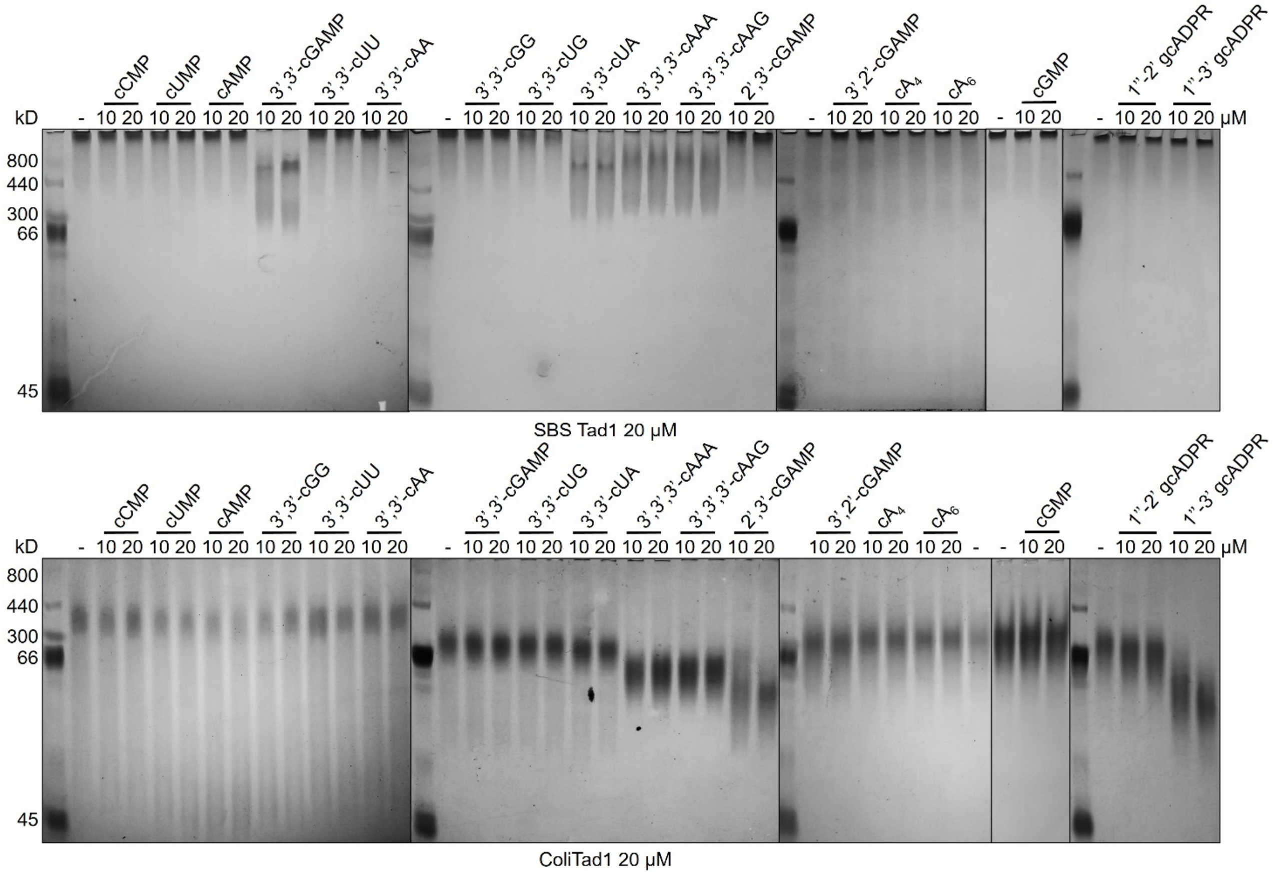


**Extended Data Figure 10. Native gel assay of SBS Tad1 and ColiTad1.**

Native PAGE showed the binding of SBS Tad1 and ColiTad1 to cyclic nucleotides.

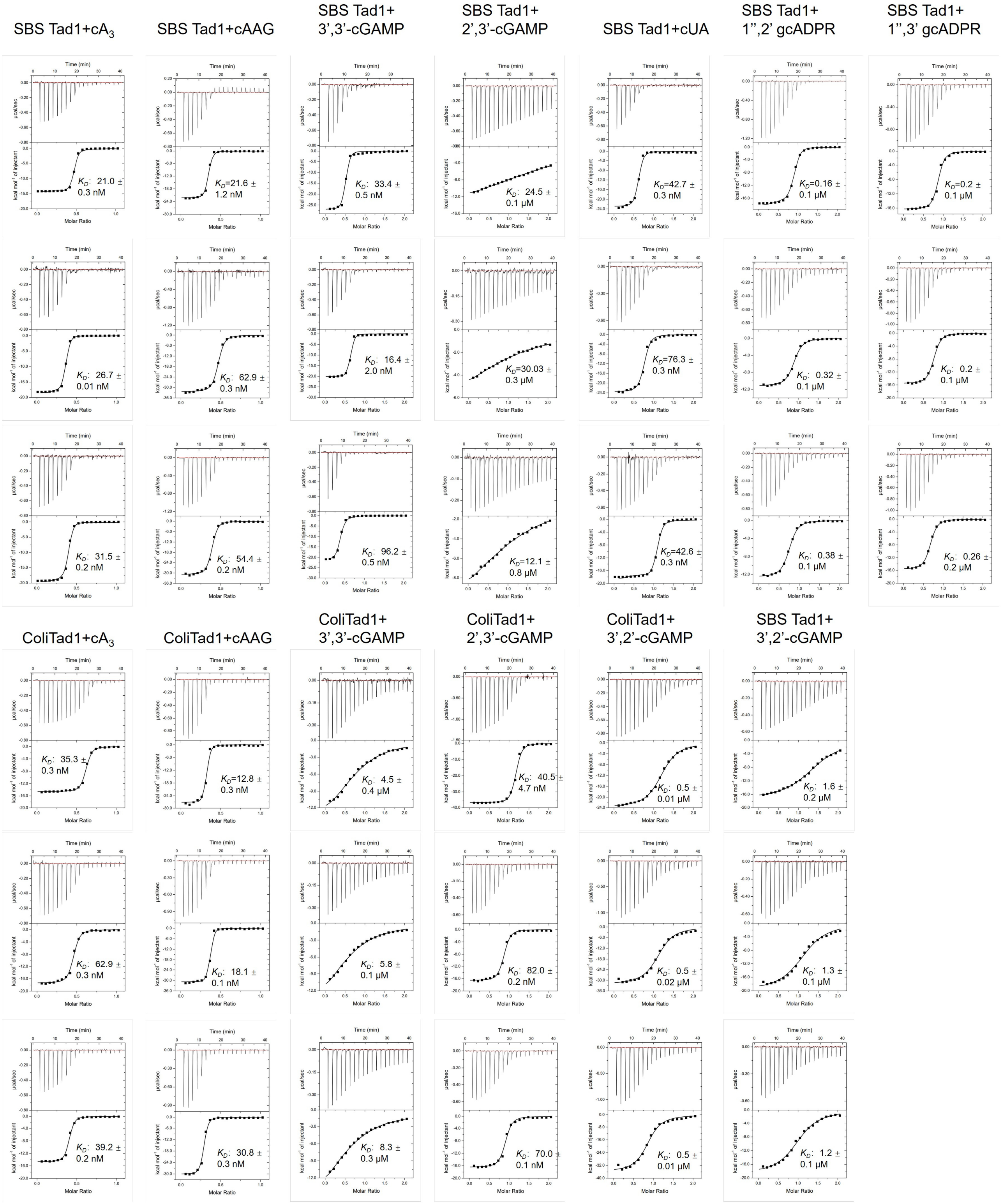


**Extended Data Figure 11. The binding spectrum of SBS Tad1 and ColiTad1 studied by ITC assay.** ITC assays to test binding of cyclic oligonucleotides to SBS Tad1 and ColiTad1.

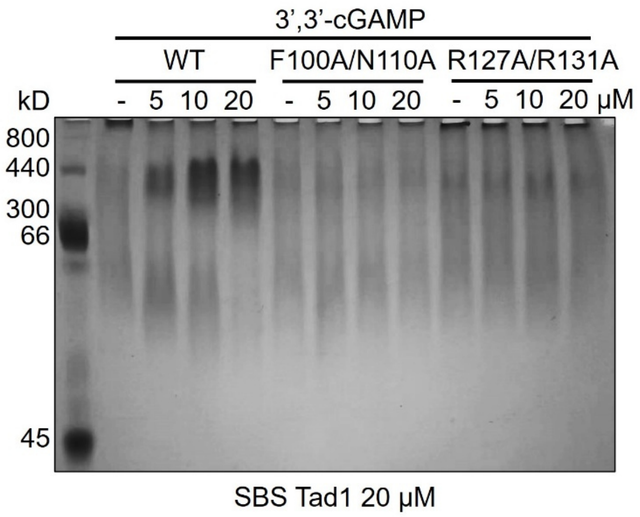


**Extended Data Figure 12. SBS Tad1 mutants lost 3',3'-cGAMP binding.**
Native PAGE showed the binding of SBSTad1 mutants to 3',3'-cGAMP.

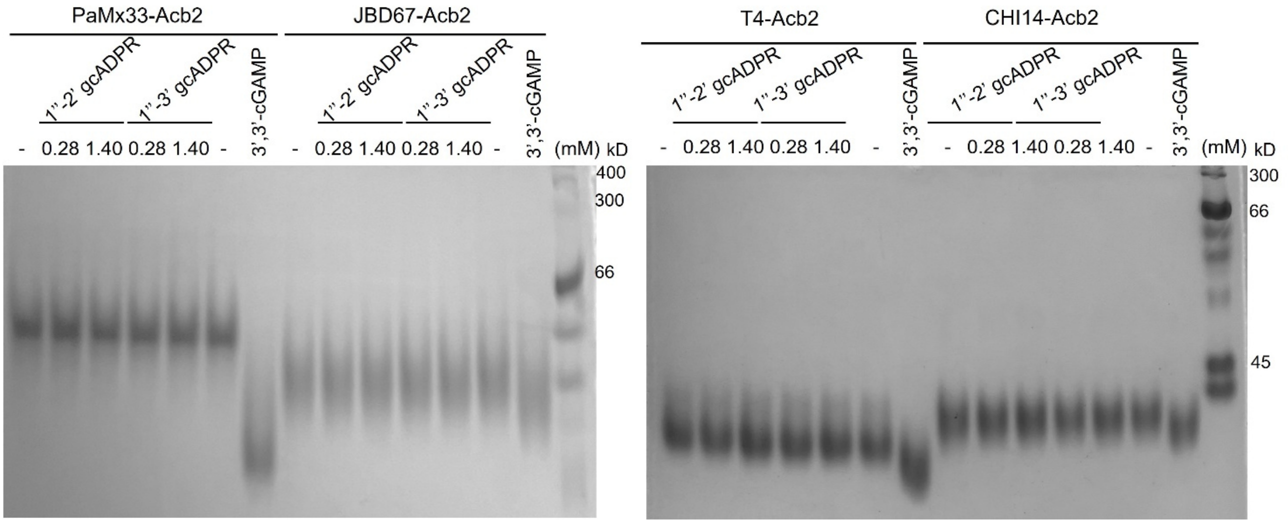


**Extended Data Figure 13. Acb2 homologs do not bind gcADPR molecules.**
All Acb2 proteins and 3',3'-cGAMP are added at a concentration of 28 µM.

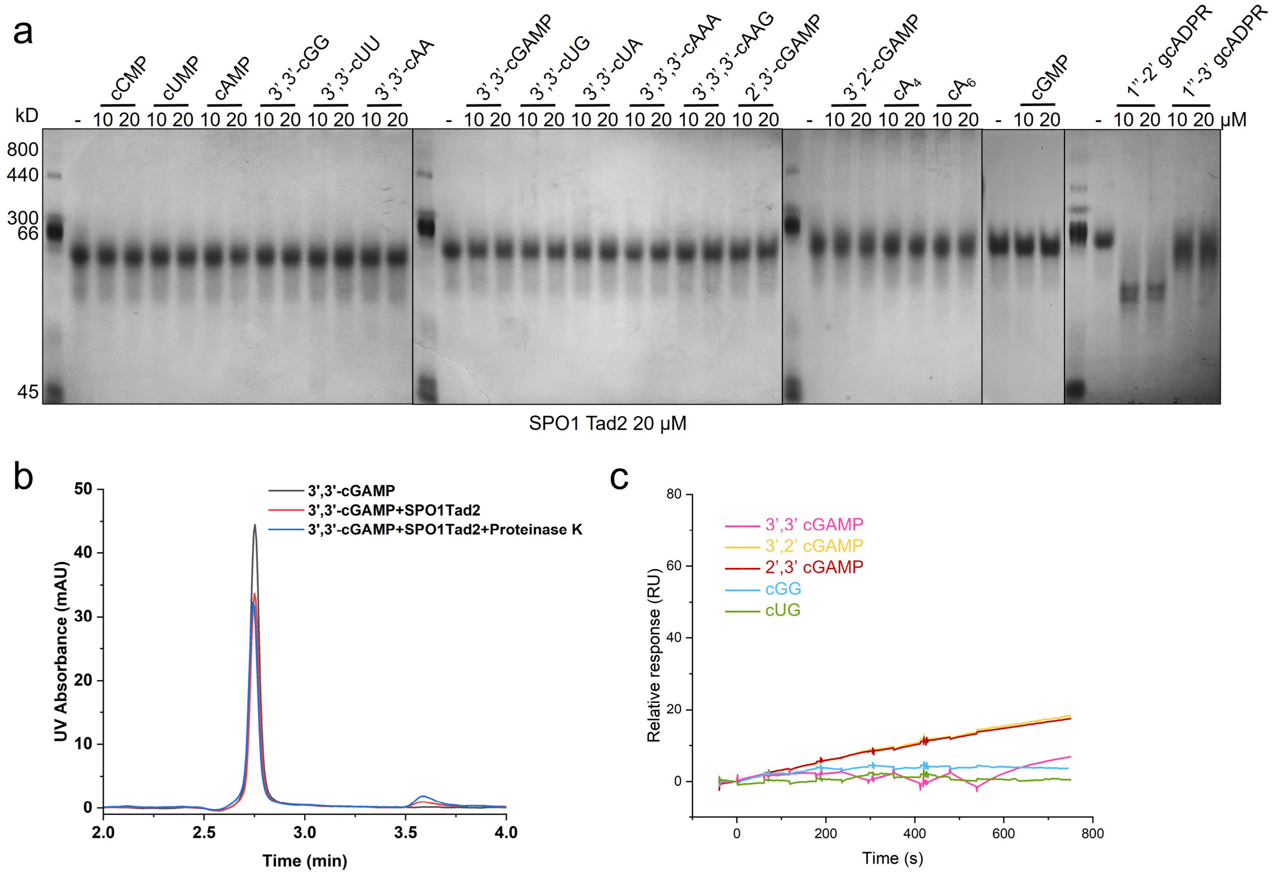


**Extended Data Figure 14. SPO1 Tad2 does not bind to cyclic dinucleotides.**

**a,** Native PAGE showed the binding of SPO1 Tad2 to cyclic nucleotides and gcADPR molecules.

**b,** The ability of SPO1 Tad2 to bind and release 3',3'-cGAMP when treated with proteinase K was analyzed by HPLC. 3',3'-cGAMP was used as a control. The remaining nucleotides after incubation with SPO1 Tad2 was tested.

**c,** Overlay of sensorgrams from surface plasmon resonance (SPR) experiments, used to determine kinetics of SPO1 Tad2 binding to cyclic dinucleotides. Data were fit with a model describing one-site binding for the ligands (black lines).

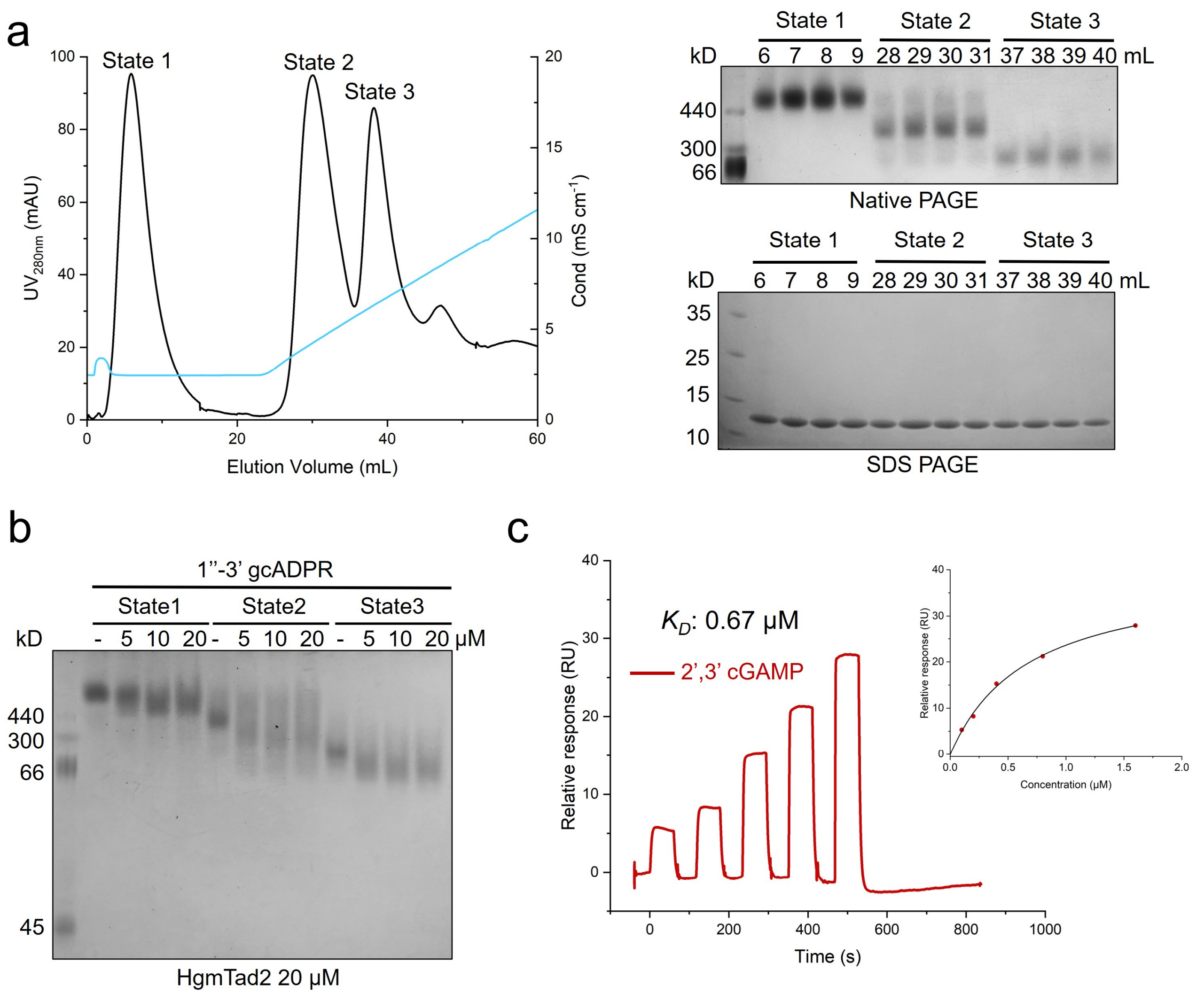


**Extended Data Figure 15. HgmTad2 binds to cGG and gcADPR molecules.**

**a,** Profile of ion exchange chromatography of HgmTad2 using Resource Q column (1 mL, GE Healthcare). Proteins in peaks 1-3 are collected separately and marked as State 1-3. The proteins in three states were then subjected to native PAGE and SDS-PAGE, respectively.

**b,** Native PAGE showed the binding of HgmTad2 in three states to 1’’,2’ gcADPR.

**c,** SPR analysis of HgmTad2 binding to 2’,3’-cGAMP. The data was fitted with affinity model and the calculated $K_D$ was shown.

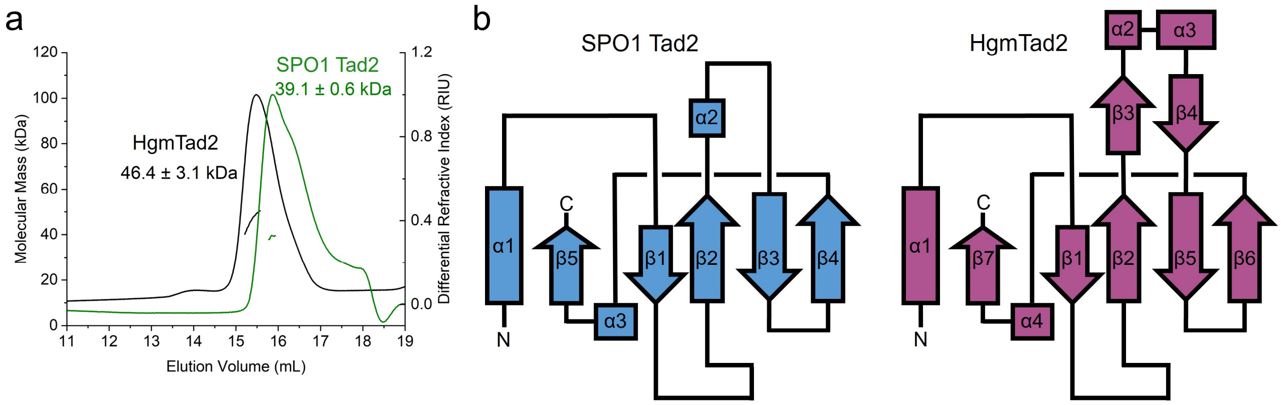


**Extended Data Figure 16. Tad2 is a tetramer.**

**a,** SLS studies of purified SPO1 Tad2 and HgmTad2. Calculated molecular weight is shown above the peaks.

**b,** Topology diagrams of SPO1 Tad2 and HgmTad2 monomers.

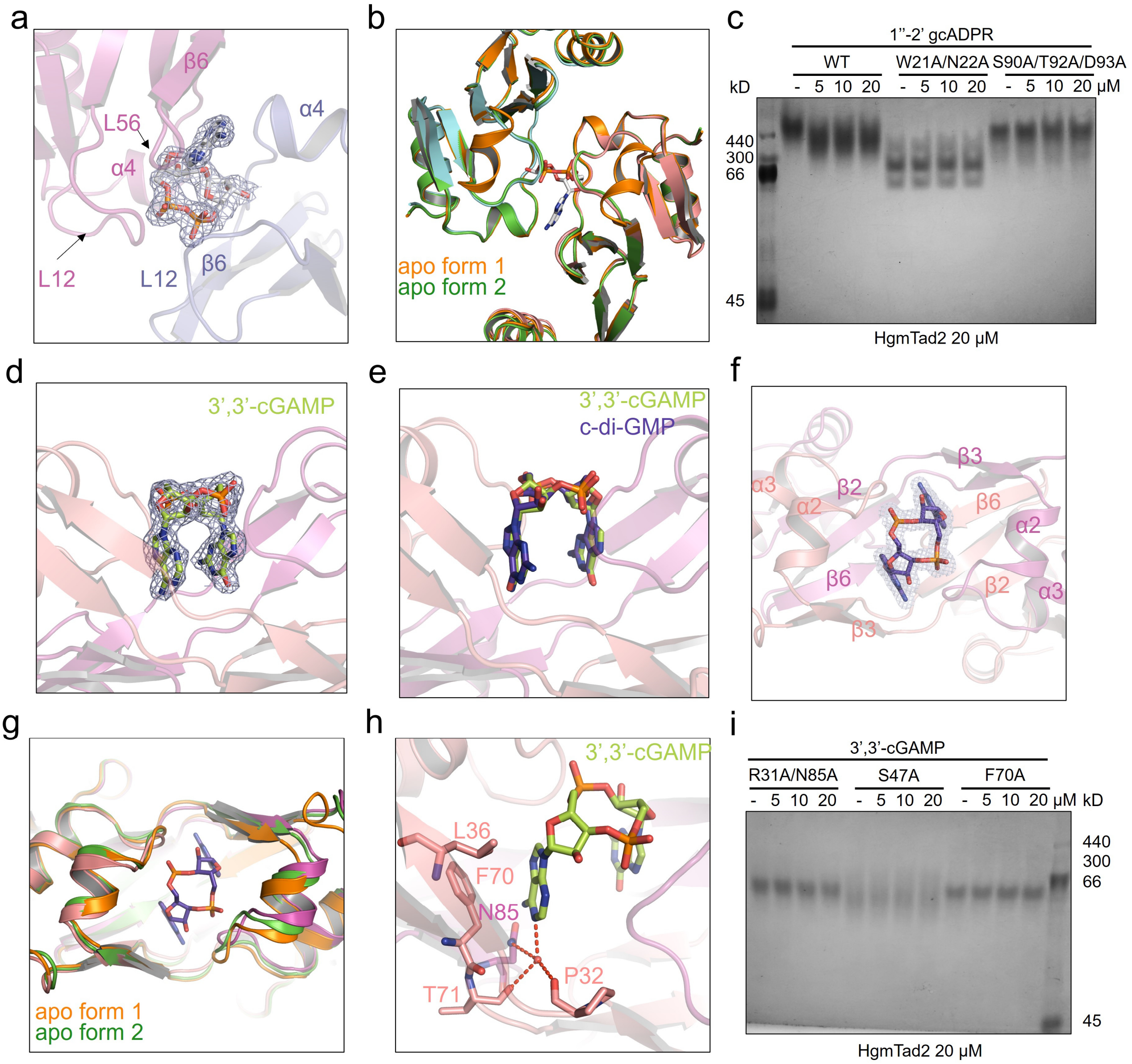

**Extended Data Figure 17. The binding pockets of 1”-2’ gcADPR and cGG.**
**a,** The binding pocket of 1”-2’ gcADPR in the HgmTad2-1”-2’ gcADPR structure. 2Fo-Fc electron density of 1”-2’ gcADPR is shown and contoured at 1 σ.
**b,** Structural superposition among HgmTad2 in the apo form (two types of conformations) and 1”-2’ gcADPR-bound form. HgmTad2 in the apo form is colored orange and green for two types of conformations, respectively. HgmTad2 in the 1”-2’ gcADPR-bound form is colored cyan and pink for the two protomers.
**c,** Native PAGE showed the binding of HgmTad2 and its mutants to 1”-2’ gcADPR.
**d,** The binding pocket of 3’,3’-cGAMP in the HgmTad2-3’,3’-cGAMP structure. 2Fo-Fc electron density of 3’,3’-cGAMP is shown and contoured at 1 σ.
**e,** Structural superposition between HgmTad2-3’,3’-cGAMP and HgmTad2-cGG. 3’,3’-cGAMP and cGG bind at the same position.
**f,** The binding pocket of cGG in the HgmTad2-cGG structure. 2Fo-Fc electron density of cGG is shown and contoured at 1 σ.
**g,** Structural superposition among HgmTad2 in the apo form (two types of conformations) and cGG-bound form. HgmTad2 in the apo form is colored orange and green for two types of conformations, respectively. HgmTad2 in the cGG-bound form is colored light magenta and pink for the two protomers.
**h,** Closer view of the binding of the adenine base of 3’,3’-cGAMP in the binding pocket of HgmTad2.
**i,** Native PAGE showed the binding of HgmTad2 mutants to 3’,3’-cGAMP.

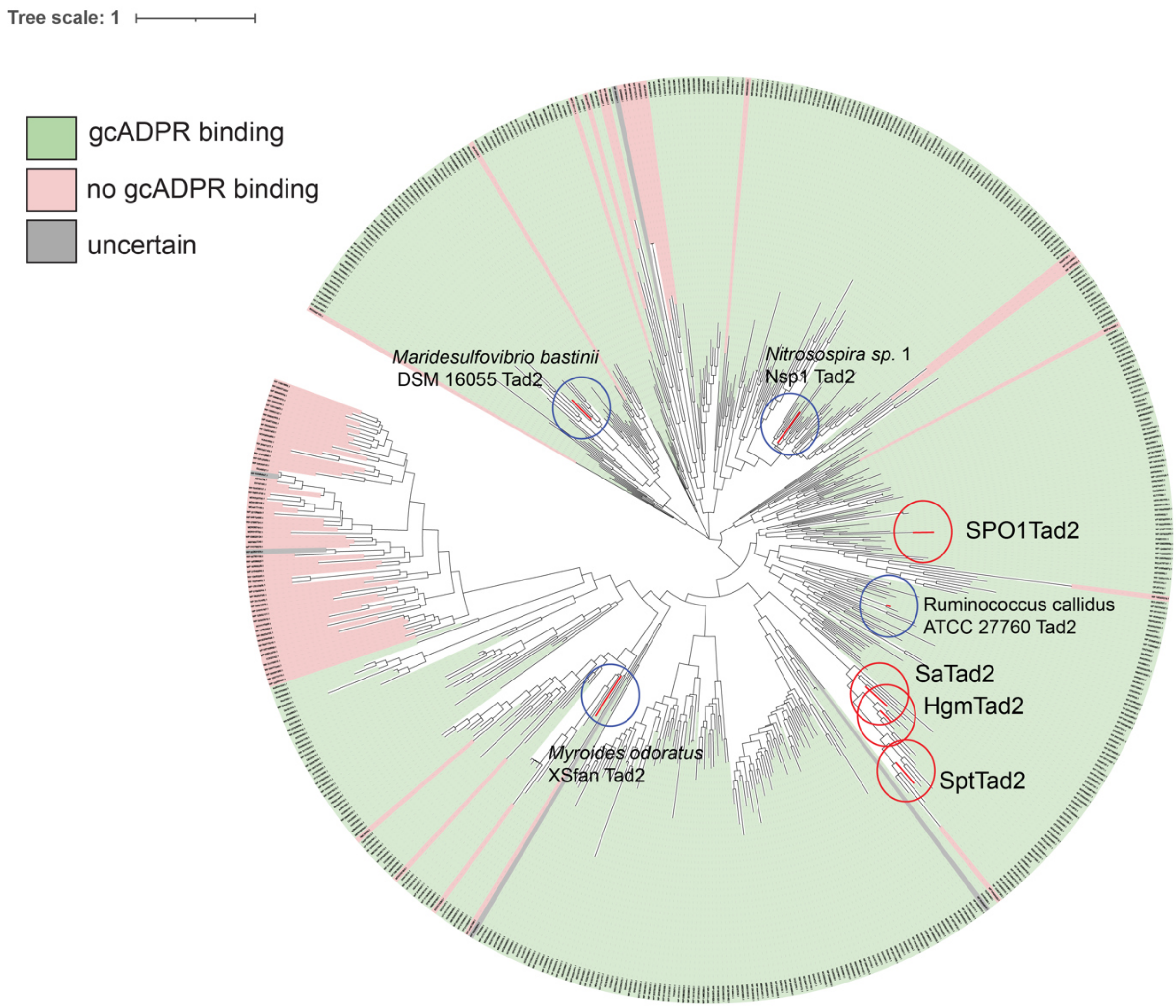


**Extended Data Figure 18. Phylogenetic analysis of Tad2 homologs.**

Phylogenetic analysis of Tad2 homologs of length 120-170 amino acids found in NCBI non-redundant protein database. The proteins gcADPR binding are predicted to be functional are indicated in green; non-functional (non-conservative substitutions in W21/N22, S90/D93 residues) - in red; mutants with non-conservative substitutions in gcADPR binding sites - in grey. Tad2 proteins used for biochemical studies are indicated on the tree by red circles. Tad2 proteins whose gcADPR binding activity was demonstrated in Yirmiya et al. 2023 are indicated by blue circles.

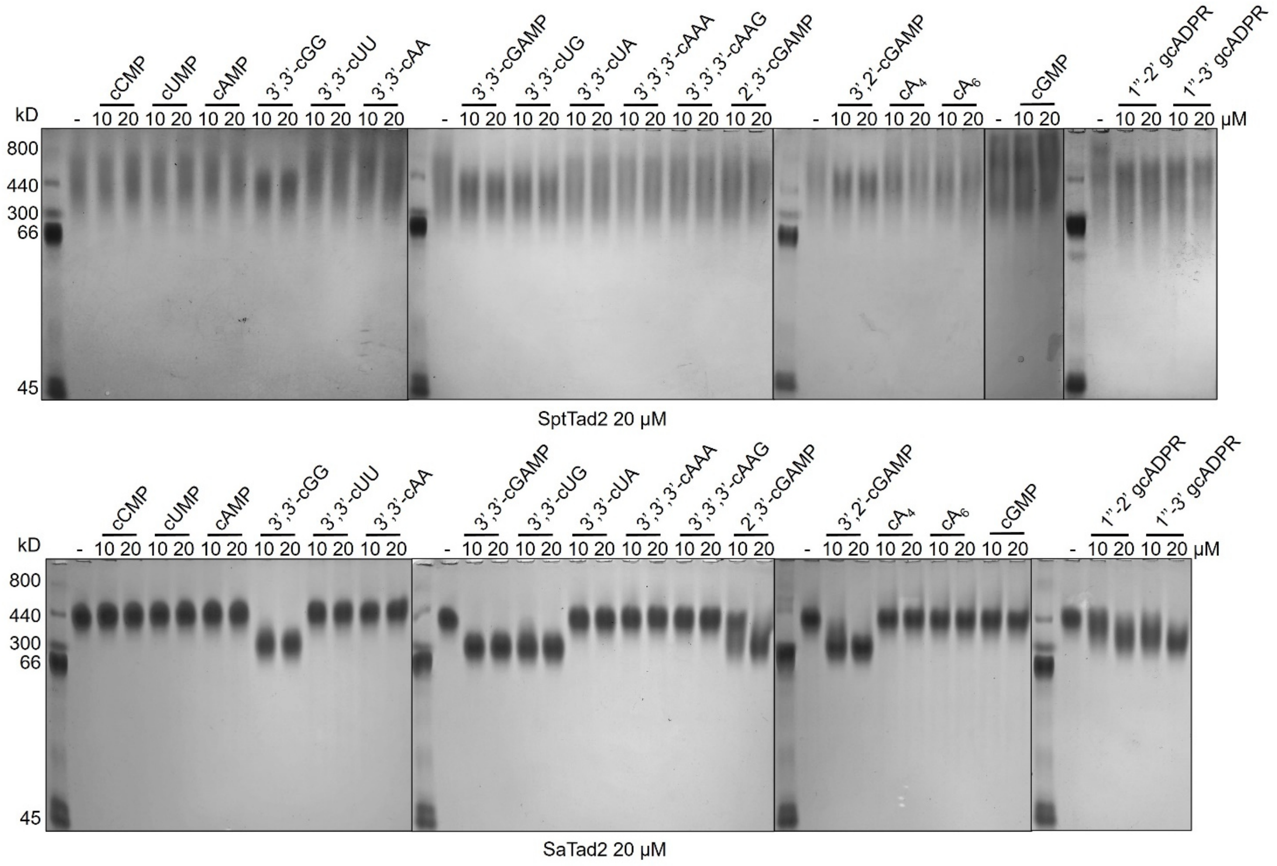


**Extended Data Figure 19. Native gel assay of SptTad2 and SaTad2.**

Native PAGE showed the binding of SptTad2 and SaTad2 to cyclic oligonucleotides and gcADPR molecules.

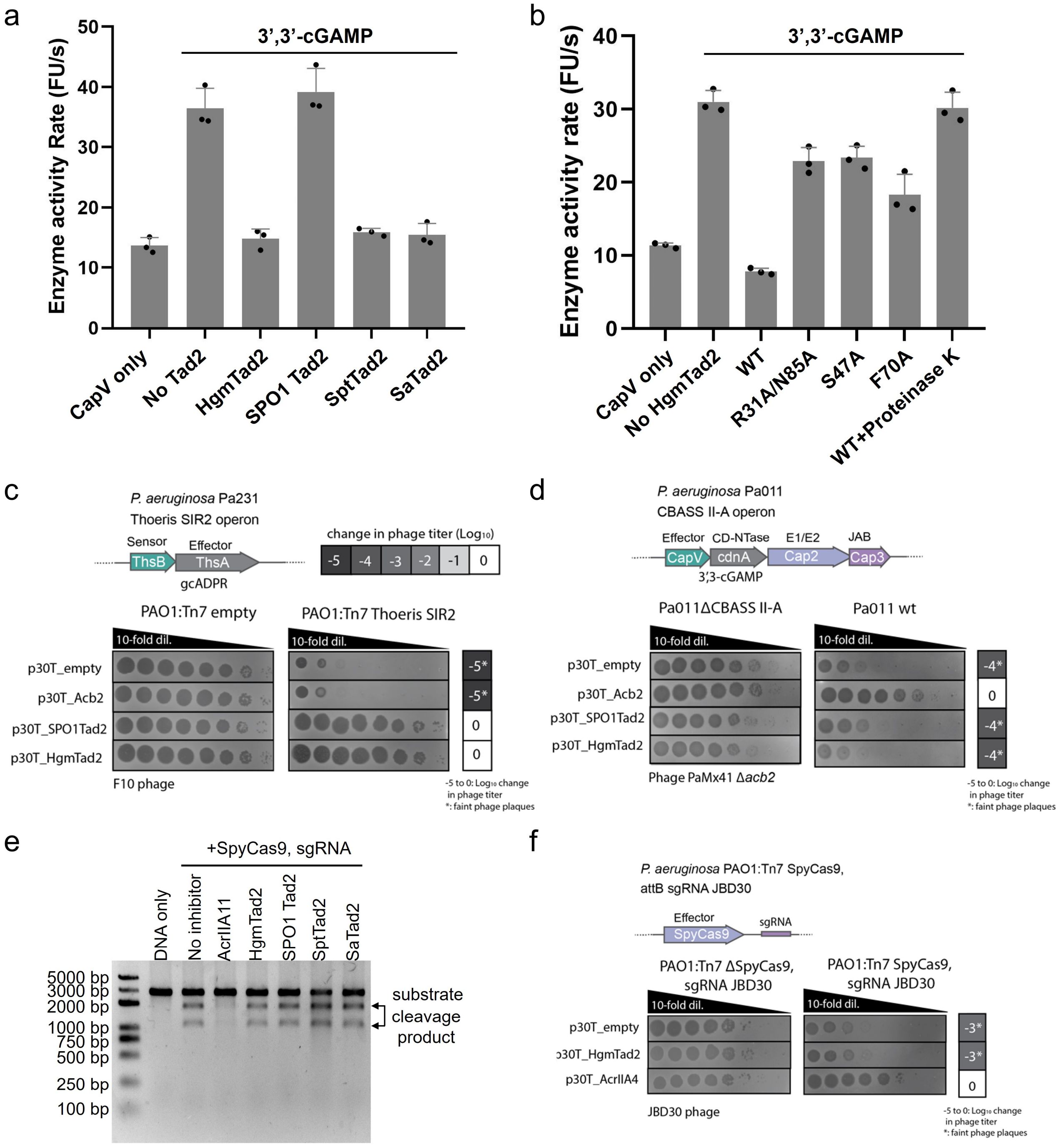


**Extended Data Figure 20. HgmTad2 has anti-Thoeris activity, but lacks anti-CBASS and anti-CRISPR-Cas activity *in vivo*.**

**a, b,** CapV enzyme activity in the presence of 3',3'-cGAMP and resorufin butyrate, which is a phospholipase substrate that emits fluorescence when hydrolyzed. The enzyme activity rate was measured by the accumulation rate of fluorescence units (FUs) per second. To test the effects of HgmTad2 or its mutants to bind and release 3',3'-cGAMP, HgmTad2, its mutants or homologs (8 μM) was incubated with 3',3'-cGAMP (0.8 μM) for 10 min and then proteinase K (0.708 mg/mL) was added to release the nucleotide from the HgmTad2 protein. Filtered nucleotide products were used for the CapV activity assay. Data are mean ± SD (n=3).

**c,** Plaque assays to test the activity of SPO1Tad2 and HgmTad2 against SIR2 containing Thoeris system *in vivo*. Organization of *P. aeruginosa* Pa231 Thoeris operon shown. F10 phage was spotted in 10-fold serial dilutions on a lawn of *P. aeruginosa* cells (PAO1) expressing Pa231 Thoeris operon genes (PAO1:Tn7 Thoeris SIR2), or cells without the system (PAO1:Tn7 empty), electroporated with pHERD30T plasmids carrying Acb2 and Tad2 genes or empty vector.

**d,** Plaque assays to test the activity of SPO1Tad2 and HgmTad2 against Type II-A CBASS *in vivo*. Organization of the *P. aeruginosa* Pa011 CBASS II-A operon shown. PaMx41Δ*acb2* was spotted in 10-fold serial dilutions on a lawn of Pa011 cells with deletion of CBASS operon (Pa011ΔCBASS II-A) or Pa011 wild type cells (Pa011 wt), electroporated with pHERD30T plasmids carrying Acb2 and Tad2 genes or empty vector.

**e,** *In vitro* DNA cleavage assay. Effect of AcrIIA11, HgmTad2 or its homologs on SpyCas9 *in vitro* cleavage activity. Presence of a cleaved DNA template indicates no inhibition of SpyCas9 activity mediated by HgmTad2 or its homologs. The concentration of SpyCas9, sgRNA, substrate DNA and AcrIIA11, HgmTad2, or its homologs is 100 nM, 150 nM, 10 nM and 10 μM, respectively.

**f,** Plaque assays to test the activity of HgmTad2 against CRISPR-Cas9 system *in vivo*. JBD30 phage was spotted in 10-fold serial dilutions on a lawn of *P. aeruginosa* cells (PAO1) expressing SpyCas9 gene and sgRNA targeting JBD30 (PAO1:Tn7 SpyCas9, sgRNA JBD30) or cells without SpyCas9 gene (PAO1:Tn7 ΔSpyCas9, sgRNA JBD30), electroporated with pHERD30T plasmids expressing HgmTad2, AcrIIA4 genes, or empty vector.

### Extended Data Table 1 Data collection and refinement statistics

| | apo-CmTad1 | CmTad1-$cA_3$ | CmTad1-cAAG | CbTad1-1",3'-gcADPR | CbTad1-1",3'-gcADPR-$cA_3$ | CbTad1-2',3'-cGAMP | CbTad1-2',3'-cGAMP-$cA_3$ | apo-HgmTad2 | HgmTad2-1",2'-gcADPR | HgmTad2-1",2'-gcADPR-cGG | HgmTad2- 1″-3′ gcADPR-cGG | HgmTad2-cGG | HgmTad2-3',3'-cGAMP | SptTad2-cGG | apo-SPO1Tad2 |
|---|---|---|---|---|---|---|---|---|---|---|---|---|---|---|---|
| **Data collection** | | | | | | | | | | | | | | | |
| Space group | $P\ 2_12_12_1$ | $P\ 2_12_12_1$ | $P\ 2_12_12_1$ | $P\ 2_13$ | $P\ 2_13$ | $P\ 2_1$ | C 2 | $P\ 3_221$ | $P\ 4_22_12$ | C 2 | $P\ 4_222$ | C 222 | $P\ 2_1$ | $P\ 6_422$ | C 2 |
| Cell dimensions | | | | | | | | | | | | | | | |
| *a*, *b*, *c* (Å) | 138.8<br>144.5<br>150.1 | 138.9<br>145.7<br>150.8 | 138.7<br>145.1<br>150.0 | 101.1<br>101.1<br>101.1 | 101.3<br>101.3<br>101.3 | 71.9<br>66.0<br>89.4 | 140.7<br>81.2<br>128.6 | 91.0<br>91.0<br>99.1 | 101.8<br>101.8<br>92.3 | 104.3<br>57.7<br>92.5 | 63.7<br>63.7<br>63.2 | 88.1<br>89.8<br>63.2 | 50.2<br>81.6<br>50.4 | 61.6<br>61.6<br>97.5 | 111.1<br>69.1<br>92.2 |
| α, β, γ (°) | 90.0<br>90.0<br>90.0 | 90.0<br>90.0<br>90.0 | 90.0<br>90.0<br>90.0 | 90.0<br>90.0<br>90.0 | 90.0<br>90.0<br>90.0 | 90.0<br>98.3<br>90.0 | 90.0<br>106.2<br>90.0 | 90.0<br>90.0<br>120.0 | 90.0<br>90.0<br>90.0 | 90.0<br>121.7<br>90.0 | 90.0<br>90.0<br>90.0 | 90.0<br>90.0<br>90.0 | 90.0<br>102.3<br>90.0 | 90.0<br>90.0<br>120.0 | 90.0<br>95.9<br>90.0 |
| Resolution (Å) | 42.28-2.56 (2.63-2.56) | 44.34-2.81 (2.88-2.81) | 50.00-3.15 (3.20-3.15) | 35.74-2.16 (2.22-2.16) | 35.80-2.31 (2.37-2.31) | 35.57-2.37 (2.43-2.37) | 35.13-1.54 (1.58-1.54) | 19.33-1.70 (1.76-1.70) | 45.56-2.10 (2.15-2.10) | 47.76-2.28 (2.34-2.28) | 36.71-1.98 (2.03-1.98) | 19.77-1.38 (1.45-1.38) | 31.40-2.11 (2.16-2.11) | 48.76-1.71 (1.75-1.71) | 35.19-2.27 (2.33-2.27) |
| $R_{sym}$ or $R_{merge}$ | 0.241 (2.489) | 0.218 (2.143) | 0.199 (133.5) | 0.070 (1.232) | 0.134 (2.305) | 0.161 (1.400) | 0.199 (1.140) | 0.277 (1.862) | 0.234 (2.055) | 0.093 (1.211) | 0.211 (2.161) | 0.107 (1.687) | 0.197 (1.259) | 0.092 (2.369) | 0.170 (2.055) |
| $I / \sigma I$ | 10.1 (1.1) | 9.4 (1.4) | 8.88 (1.0) | 31.3 (2.3) | 22.0 (2.3) | 10.2 (1.8) | 5.8 (1.6) | 7.8 (2.6) | 15.7 (5.8) | 11.5 (1.7) | 18.6 (8.1) | 15.0 (2.0) | 8.5 (2.0) | 25.8 (2.1) | 8.9 (1.6) |
| Completeness (%) | 100 (100) | 100 (100) | 99.2 (97.8) | 100 (100) | 100 (100) | 99.9 (100) | 100 (99.9) | 100 (100) | 100 (100) | 98.2 (97.2) | 100 (100) | 97.9 (95.8) | 99.7 (99.7) | 99.2 (99.7) | 99.3 (98.5) |
| Redundancy | 12.9 (12.8) | 13.1 (13.7) | 5.0 (4.1) | 35.1 (17.8) | 40.0 (41.1) | 6.8 (7.1) | 6.1 (4.8) | 39.9 (40.0) | 25.3 (24.8) | 6.8 (6.3) | 24.3 (26.0) | 26.4 (27.6) | 6.5 (6.5) | 35.3 (32.4) | 6.6 (6.8) |
| **Refinement** | | | | | | | | | | | | | | | |
| Resolution (Å) | 41.13-2.56 (2.65-2.56) | 44.14-2.80 (2.91-2.80) | 34.76-3.15 (3.26-3.15) | 31.97-2.16 (2.24-2.16) | 32.02-2.31 (2.39-2.31) | 31.47-2.37 (2.46-2.37) | 38.84-1.54 (1.60-1.54) | 19.33-1.70 (1.76-1.70) | 44.58-2.10 (2.18-2.10) | 26.97-2.28 (2.36-2.28) | 31.86-1.98 (2.05-1.98) | 19.77-1.38 (1.43-1.38) | 31.36-2.11 (2.19-2.11) | 30.77-1.71 (1.77-1.71) | 32.51-2.27 (2.35-2.27) |

| | | | | | | | | | | | | | | | |
|---|---|---|---|---|---|---|---|---|---|---|---|---|---|---|---|
| No. reflections | 97006 (9462) | 74744 (6907) | 52431 (4950) | 18756 (1878) | 15410 (1500) | 33876 (3364) | 204857 (20288) | 51811 (5206) | 28922 (2847) | 21044 (2047) | 9542 (932) | 50735 (4912) | 22794 (2306) | 12276 (1187) | 31937 (3147) |
| $R_{work}$ / $R_{free}$ | 0.229/ 0.262 | 0.227/ 0.262 | 0.202/ 0.239 | 0.205/ 0.250 | 0.204/ 0.250 | 0.195/ 0.241 | 0.239/ 0.262 | 0.213/ 0.228 | 0.180/ 0.222 | 0.228/ 0.289 | 0.200/ 0.238 | 0.203/ 0.218 | 0.233/ 0.272 | 0.201/ 0.236 | 0.234/ 0.261 |
| No. atoms | 13140 | 12833 | 12885 | 2175 | 2303 | 6369 | 10772 | 3659 | 3622 | 3546 | 985 | 2001 | 3451 | 907 | 4260 |
| Protein | 12583 | 12567 | 12615 | 2059 | 2048 | 5982 | 8994 | 3288 | 3254 | 3288 | 827 | 1644 | 3288 | 816 | 4189 |
| Ligand/ion | 2 | 266 | 270 | 70 | 202 | 270 | 603 | / | 140 | 232 | 81 | 92 | 90 | 46 | / |
| Water | 555 | / | / | 45 | 53 | 117 | 1175 | 371 | 228 | 26 | 77 | 265 | 73 | 45 | 71 |
| *B*-factors | 48.59 | 63.41 | 73.72 | 60.66 | 59.78 | 56.45 | 17.60 | 35.16 | 30.35 | 61.49 | 27.59 | 25.91 | 46.47 | 40.19 | 50.17 |
| Protein | 48.86 | 63.51 | 73.97 | 60.78 | 60.68 | 56.73 | 17.17 | 34.44 | 30.50 | 61.78 | 27.09 | 24.52 | 46.55 | 40.67 | 50.28 |
| Ligand/ion | 55.23 | 58.84 | 62.16 | 57.73 | 52.81 | 53.66 | 11.78 | / | 19.46 | 58.26 | 25.75 | 20.00 | 45.34 | 27.52 | / |
| Water | 42.40 | / | / | 56.78 | 51.52 | 48.72 | 23.83 | 41.50 | 34.96 | 52.75 | 34.90 | 36.59 | 44.33 | 44.39 | 43.49 |
| R.m.s. deviations | | | | | | | | | | | | | | | |
| Bond lengths (Å) | 0.011 | 0.011 | 0.011 | 0.006 | 0.010 | 0.009 | 0.009 | 0.013 | 0.010 | 0.010 | 0.014 | 0.013 | 0.01 | 0.012 | 0.017 |
| Bond angles (°) | 1.18 | 1.24 | 1.26 | 1.02 | 1.43 | 1.17 | 1.24 | 1.52 | 1.09 | 1.18 | 1.54 | 1.42 | 1.13 | 1.30 | 1.59 |

For each structure one crystal was used.

*Values in parentheses are for highest-resolution shell.
